# Mapping non-coding functional elements in allotetraploid *Cyprinus carpio* embryo development reveals subgenome variation of transcription regulation

**DOI:** 10.64898/2026.06.22.733679

**Authors:** Ada Jimenez-Gonzalez, Alba Madrero Pardo, Yavor Hadzhiev, Annemiek Blasweiler, Bojan Zunar, Zsolt Csenki-Bakos, Tamás Müller, Hendrik-Jan Megens, Geert Frits Wiegertjes, Boris Lenhard, Damir Baranasic, Ferenc Müller

## Abstract

Common carp (*Cyprinus carpio*) is an important freshwater species for ornamental and aquaculture purposes, and a key cyprinid model for studying allotetraploidy. Its two chromosomally-separated subgenomes show distinct gene expression profiles, but how their regulatory landscapes control gene expression dynamics during development remains unknown. We generated a regulatory atlas by combining transcriptomes across 12 developmental stages with chromatin accessibility maps, transcription start sites and gene regulation-associated histone post-translational modifications. Subgenome-specific annotation and comparison of 254,276 developmental regulatory elements (PADREs) revealed that regulatory subgenome divergence is most prominent during early development, converging toward the phylotypic period, mirroring expression convergence between subgenomes at the same stages. This dynamic was driven by enhancers, while promoters maintained a more stable subgenome bias, extending the hourglass model of developmental constraint to allotetraploid subgenome regulation. Subgenome-specific enhancers were preferentially retained in subgenome B, whereas subgenome A shifted toward homeologous enhancer activity near the phylotypic stage, indicating directional regulatory divergence between subgenomes. Comparison with zebrafish revealed high concordance with sequence conservation and that subgenome B retained more ancestral cyprinid regulatory elements than subgenome A. This developmental regulatory atlas provides a foundational resource for investigating cis-regulatory evolution following the fourth round of vertebrate genome duplication.

## Introduction

The common carp (*Cyprinus carpio*) is one of the most studied non-model freshwater teleosts, owing to its ecological, aquaculture, and cultural significance. Ohno first proposed carp as tetraploid species and leveraged it as a key example for his hypothesis on genome duplication as major driver of evolutionary change^1^. Genome duplication is often argued to underlie common carp’s remarkable environmental adaptability, making it a valuable genetic model for adaptive mechanisms, including immune responses that improve disease resistance and tolerance of hypoxia and extreme temperatures^2–4^. How the evolution of its two subgenomes contributes to such physiological and ecological adaptation, remains poorly understood.

Despite the rarity of polyploidisation among vertebrates, the subfamily Cyprininae, which includes common carp and the tetraploid goldfish, is the largest polyploid vertebrate group, with over 600 species^5^. The carp genome is small for a tetraploid (1.6 Gb) and arose by allotetraploidisation during a fourth round of genome duplication (4R)^6^. Hybridisation occured between a Barbinae species and a now-extinct unidentified relative approximately 11 to 13 million years ago (Myr; the progenitors estimated to be ∼23 Myr apart) and gave rise to two subgenomes, which have remained chromosomally distinct with largely intact homeologous chromosomes and a broadly balanced gene number^7^.

Functional divergence between the two subgenomes during ontogeny has been studied to explain polyploid complexity, adaptation, and phenotypic traits. Subgenome expression dominance, dosage compensation, and dynamic functionalisation of homeologs have all been described ^7–9^. These biases are largely category-specific: subgenome A dominates genes for nucleobase biosynthesis and metabolism, whereas subgenome B-biased genes are enriched in redox processes, stress responses, and DNA repair^7^. Hypoxia tolerance has been linked to subgenome A genes for macromolecule biosynthesis and protein binding, while subgenome B bias has been implicated in the regulation of signalling, responses to stimuli, and cell communication ^8^, while no strong evidence of subgenome dominance was found for antiviral responses^10^. What remains unknown is how these genes are regulated, and which cis-regulatory elements drive expression bias and divergence in this allotetraploid teleost.

Notably, regulatory divergence in the allotetraploid frog *Xenopus laevis* was shown to underlie subgenome-specific gene expression during the maternal-to-zygotic transition^11^. Among vertebrates, developmental transcriptomes are most constrained at the mid-embryonic phylotypic stage, resembling an hourglass pattern of evolutionary conservation ^12,13^, yet whether this constraint is reflected in regulatory dynamic and whether it relates to the divergence between a polyploid’s subgenomes is unknown. Addressing these questions, and supporting the use of common carp in developmental and aquaculture research, requires functional annotation of the carp genome, including mapping of its cis-regulatory elements (CREs), such as enhancers and promoters. A regulatory atlas will allow CRE evolution to be traced and CRE activity to be linked to transcriptional variation between homeologs through development. Carp and zebrafish (*Danio rerio*), a widely used genetic model with a well-annotated non-coding genome^14^, are both cyprinids separated by 42 to 90 Myr^7,15^; the high conservation of predicted regulatory elements among cyprinids^16^ makes zebrafish a powerful outgroup comparator for a non-model tetraploid of high ecological, evolutionary, developmental and aquaculture value.

## Results

### A multiomic catalogue of the functional elements of carp embryo development

In this study, we generated the first genome-wide annotation map of cis-regulatory elements during common carp ontogeny and provided the foundation for explaining transcriptional activity of an allotetraploid teleost (**Figure 1a**). We assembled a comprehensive collection of multiomic datasets spanning embryo development and comprising 119 sequencing libraries. Specifically, we generated 36 transcriptomes starting from unfertilised eggs through maternal to zygotic transition, gastrulation, neurulation and organogenesis. Embryonic stages were identified based on the remarkably high anatomical similarity with zebrafish^17^ (**Supplementary Figure 1**, **Supplementary Table 1**). We complemented the transcriptomes with high resolution promoter identification by transcription start site mapping using Cap Analysis of Gene Expression (CAGE)-seq. In addition, we mapped cis-regulatory element-associated chromatin features such as chromatin accessibility by ATAC-seq and histone post-translational modifications by ChIP-seq targeting gene activation (H3K4me3 and H3K27ac) and repression-associated (H3K27me3) histone post-translational modification marks. Embryo samples were obtained from the strain with the latest genome assembly available in Ensembl (Cypcar_WagV4. 0) (**Figure 1b**). Standardised metadata, and standard operating protocols (SOPs) of genomics libraries (**Supplementary Tables 2-5** and FAANG dataportal) followed the guidelines by the Functional Annotation of Animal Genomes (FAANG) initiative^18^ and AQUA-FAANG consortium^19^.

**Figure 1.**
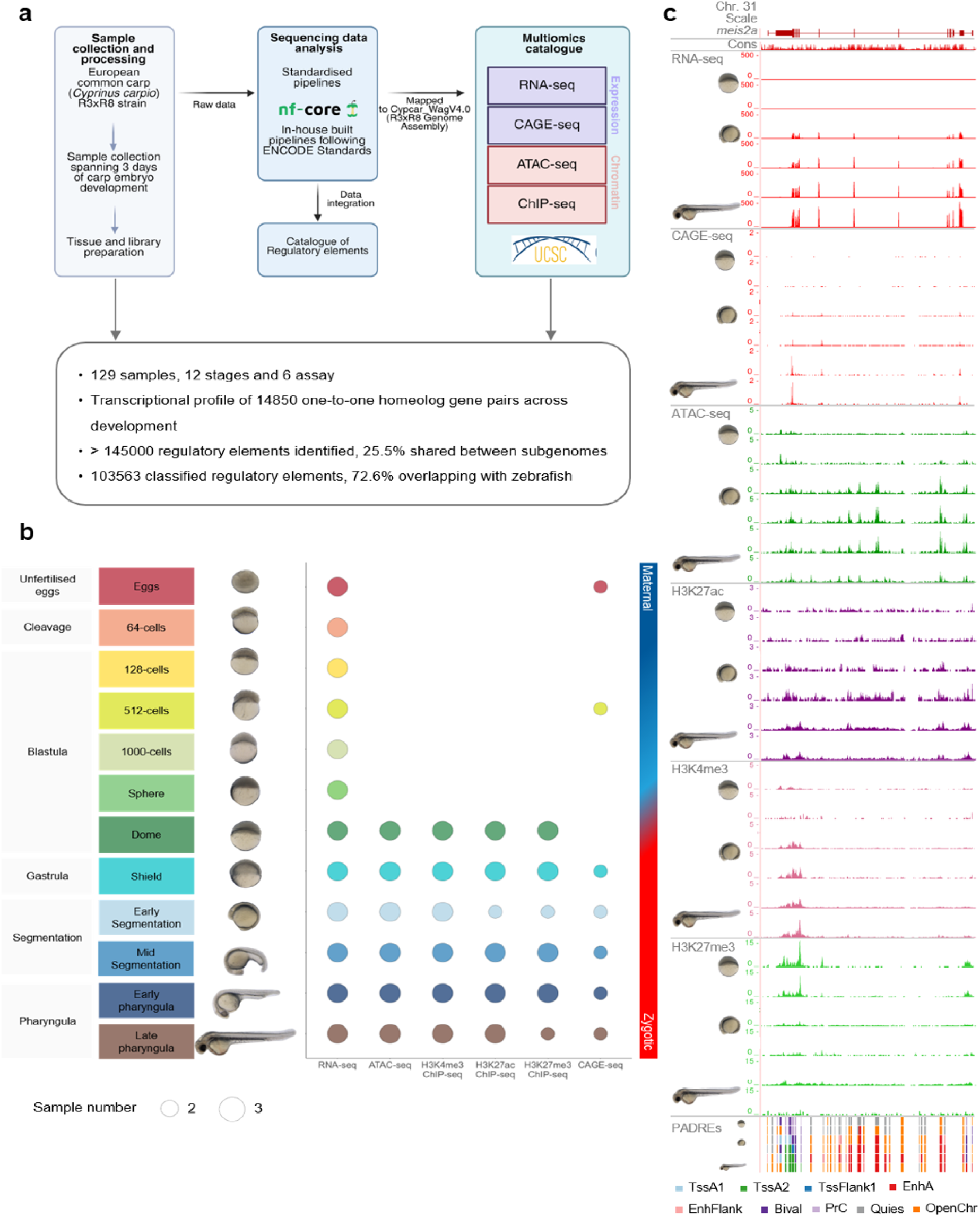
Collection and annotation of carp developmental multiomic data. **a,** Pipeline of data generation and analysis. **b,** Stages of development for which genomics data have been produced in duplicates or triplicates. **c,** Example screenshots of the *meis2* locus at the UCSC genome browser and example developmental stages for genomics data used for regulatory element annotation and gene expression analyses. Cons: phastCons track.

We organised the processed datasets into track hubs for the UCSC Genome Browser (**Figure 1c**). These track hubs (link: https://genome-euro.ucsc.edu/s/ajimglez/Manuscript_session) are accompanied by information on annotation of candidate CREs during common carp development. Together, we identified 45107 expressed genes and defined a set of 14849 one-to-one homeolog gene pairs to investigate transcriptional subgenome divergence and we annotated 255,028 regulatory elements of which 254,276 were successfully assigned to subgenomes.

### Developmental dynamics and functional divergence of gene expression asymmetry in subgenomes

We characterised the dynamic changes in transcriptomes during carp ontogeny, leveraging the temporal resolution in our transcriptomic datasets spanning development from unfertilised egg to late pharyngula. This catalogue expands on existing transcriptome compendia of carp development^20^, increases temporal resolution thus providing a unique opportunity to focus on subgenome dynamics at major developmental events such as the maternal-to-zygotic transition (MZT) and offers linking transcriptional programmes with dynamics of regulatory architecture.

Reproducibility was supported by principal component analysis, which revealed clustering of replicates from the same stage with PC1 (64.14% variance explained) and sample correlation analysis (**Supplementary Figure 2a**) indicating sample separation by developmental stage and reflecting major transcriptional transitions, including the MZT (**Fig. 2a**). Gene expression levels increased progressively across stages, with late pharyngula embryos showing the highest degree of transcriptional activity (**Supplementary Figure 2b**). By ranking genes by their developmental stage of maximum expression across each stage replicate, we identified stage-specific expression programmes (**Figure 2b**,).

**Figure 2.**
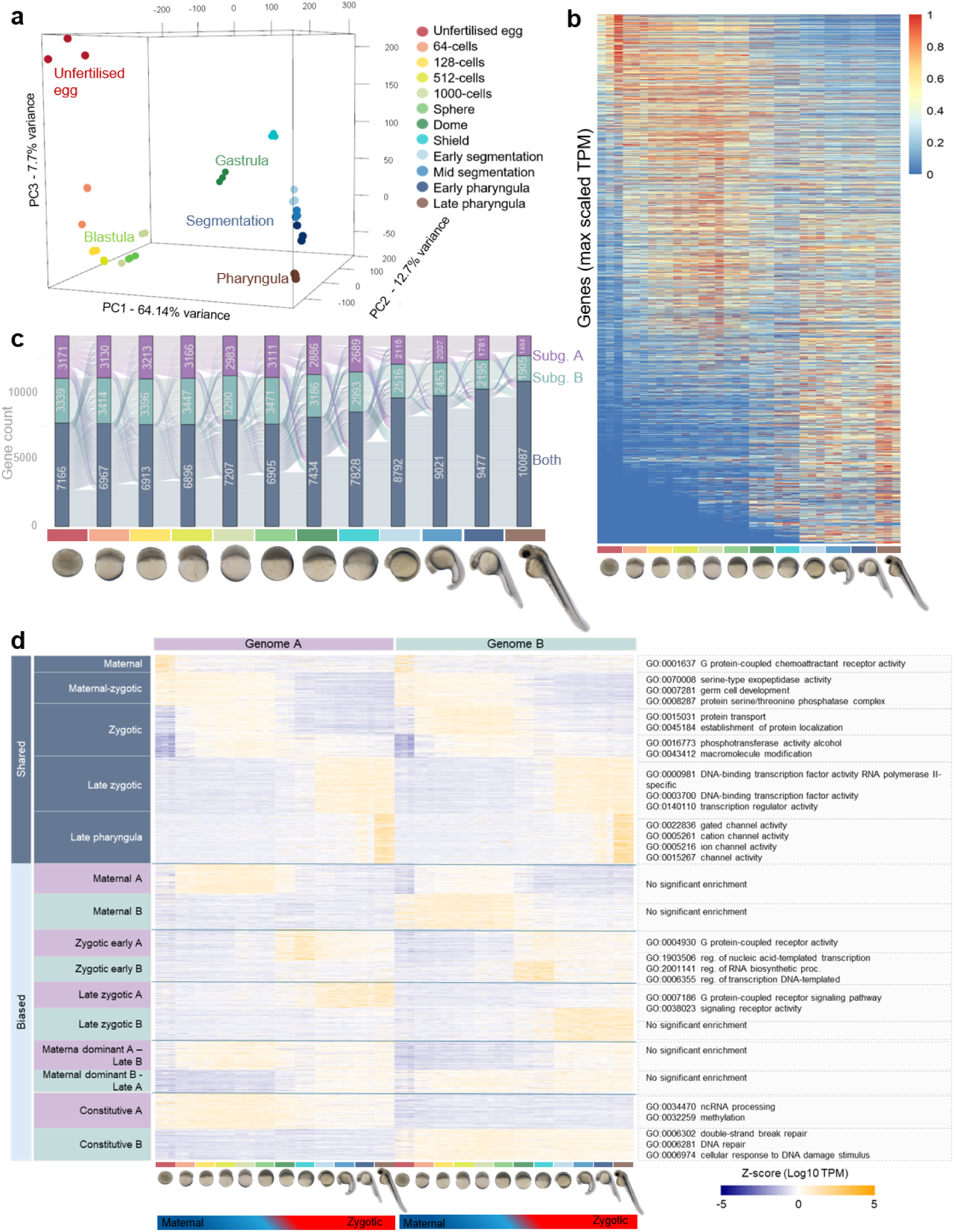
Subgenome-associated distinction of developmental dynamics of the carp transcriptome. **a,** Principal Component Analysis of gene expression at the stages as indicated. **b,** Heatmap of gene expression during development for genes with TPM>0. TPM values were scaled to the maximum expression for each gene across all stages and ranked to the highest expression stage. **c,** Alluvial plot of gene expression dynamics of 14849 homeolog gene pairs between subgenomes in development. **d**, Heatmap and clustering by stages of gene expression of homeolog gene pairs. GO: Gene ontology terms of biological process enriched for subgenome-specific clusters.

To assess changes in gene expression across development, we applied a likelihood ratio test and found that 36054 out of the 38114 expressed genes were differentially expressed across stages (>10 counts) (**Supplementary Figure 3a**, **Supplementary Table 6**). In contrast to previous transcriptome studies, we noticed changes at very early development between fertilised eggs and 64-cell stage embryos. Differential expression analysis between these stages revealed 4453 upregulated genes in the egg representing maternal mRNAs, which degraded by the 64 cell stage. Furthermore, 2931 upregulated genes in 64-cell embryos (**Supplementary Figure 3b, Supplementary Table 7**) were found, which were likely maternal mRNAs similarly subjected to delayed polyadenylation during early stages of development as was described in zebrafish^21^. Our analysis comparing the genes undergoing delayed polyadenylation in zebrafish and the upregulated genes in the 64-cell stage in carp showed an overlap of 879 genes between these two sets (17.4%, Fisher’s exact test, p=2^−16^). Gene ontology analysis of this subset revealed similar biological processes to that in zebrafish, including protein transport, protein localisation, and cellular macromolecular metabolic process (**Supplementary Figure 4**).

We next examined the transcriptome dynamics between the two carp subgenomes. To compare gene expression directly, we identified one-to-one homeolog genes resolving 14849 pairs (**Figure 2c, Supplementary Table 8**). We used Ensembl gene IDs as an anchor to split the counts assigned to the transcripts from homeologs. Across development, matching expression of both homeologs was more prevalent than subgenome-bias, reaching the highest frequency of matching pairs at the late pharyngula stage. In contrast, the earliest stages were characterised by higher transcriptional asymmetry, with homeologs from subgenome B being more expressed than homeologs from subgenome A (**Supplementary Figure 5a,b**). To investigate the functional relevance of such transcriptional patterns, we applied K-means clustering to identify genes, which were either subgenome-shared or biased during development (**Figure 2d, Supplementary Figure 6, Supplementary Table 9**). We further classified the genes based on their developmental pattern of expression. We grouped co-expressed homeologs into five temporal classes: maternal, maternal-zygotic, zygotic, late zygotic, and late pharyngula. Subgenome-biased genes were grouped into 10 categories, with 3 temporal classes split by subgenome (maternal-zygotic, ZGA, and late zygotic), each assigned to subgenome A or B. Four constitutive patterns captured cross-subgenome specificity: constitutive in A only, constitutive in B only, constitutive in A with late onset in B, and constitutive in B with late onset in A. Gene Ontology (GO) enrichment analysis revealed that symmetrically expressed maternal and maternal–zygotic genes were enriched for functions related to germ cell development and protein serine/threonine phosphatase complexes. In contrast, no significant GO enrichment was detected for subgenome-biased genes at these early stages. However, ZGA-associated genes preferentially expressed from subgenome A were enriched for G protein–coupled receptor activity, whereas those expressed from subgenome B were enriched for transcription-related functional categories.

### Cis-regulatory element identification and functional classification by chromatin segmentation

Next, we utilised our chromatin feature detection to characterise the regulatory landscape during common carp development from dome to late pharyngula (**Figure 1, Supplementary Figure 7,8**). We used H3K4me3 (active or activation-ready promoters), H3K27ac (active promoters and enhancers), and H3K27me3 (associated with Polycomb-mediated gene repression) marks to categorise regulatory element dynamics. Following an approach used in the zebrafish regulatory element atlas^14^ we integrated the above histone marks with ATAC-seq using ChromHMM^22^ and segmented the accessible chromatin into ten functional states (**Figure 3a, Supplementary Table 11**) which we called Predicted ATAC-supported Developmental Regulatory Elements (PADREs). These PADRES were classified into common regulatory element classes, including active promoters (TssA1, A2), flanking promoter regions (TssFlank1,2), active enhancers (EnhA), enhancer-flanking regions (EnhFlank), open chromatin lacking histone marks (OpenChr), Polycomb-repressed regions (PrC), candidate bivalent domains co-marked by H3K4me3/H3K27ac H3K27me3 (Bival), and quiescent chromatin (Quies) (**Figure 3a,b**). Chromatin segmentation distinguished two promoter classes: canonical promoters marked by both H3K4me3 and H3K27ac (TssA1) and non-canonical promoters carrying H3K4me3 but lacking H3K27ac (TssA2), potentially reflecting alternative or poised promoter states (**Figure 3b**).

**Figure 3.**
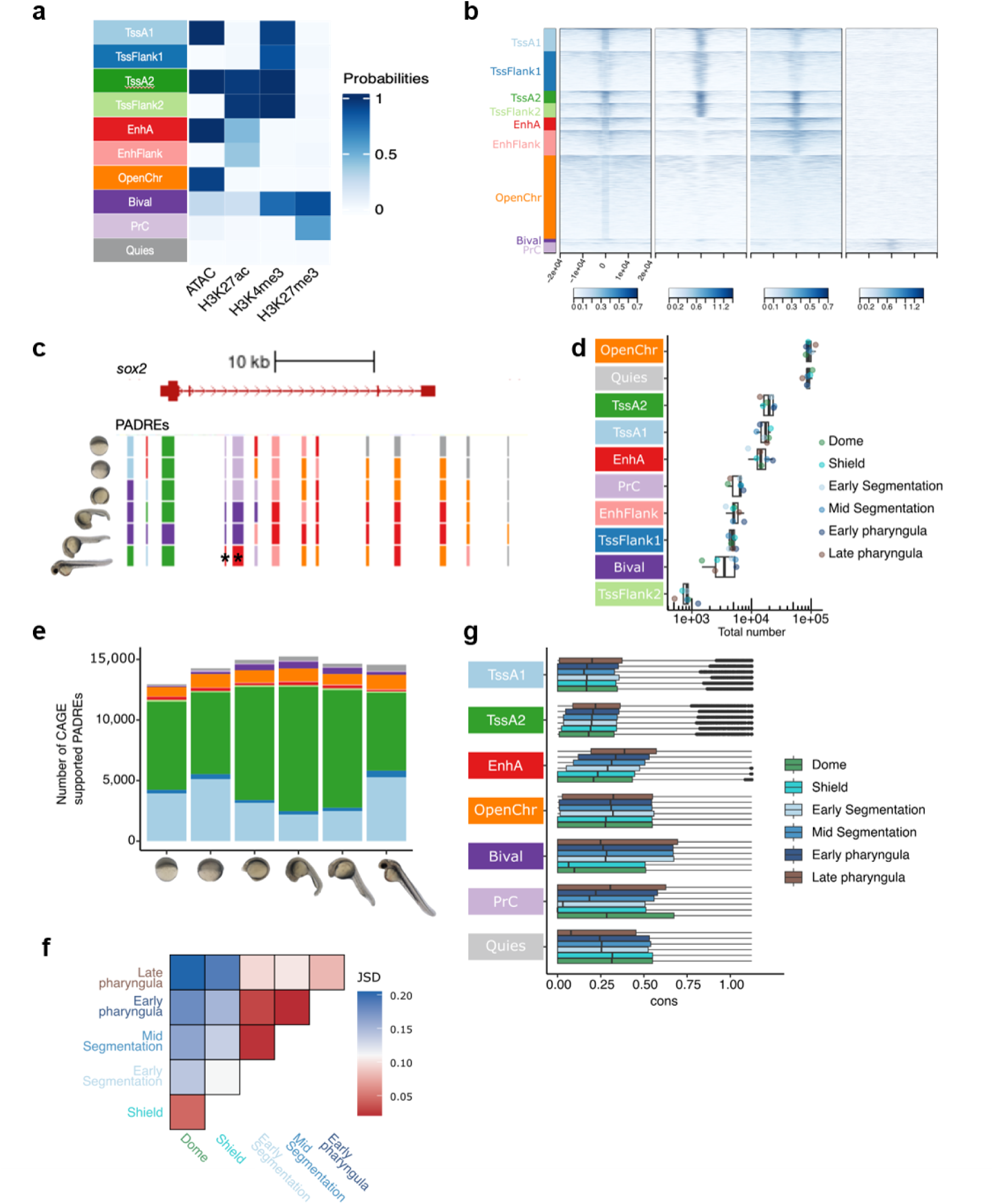
Annotation and Classification of cis-regulatory elements in carp development. **a,** Occurrence probabilities of chromatin marks for ChromHMM states and that of ATAC open chromatin. The states function was manually assigned using The Roadmap Epigenomic annotations. TssA, Active TSS, TssFlank,TSS flanking region, EnhA, active enhancer, EnhFlank, enhancer flanking region, PrC, polycomb repressed regions, Quies, quiescent state. **b**, Aggregation heat map of the genomic features as indicated. Signals aligned to peak summit for loci of ChromHMM classification. **c,** Genome browser screenshot of *sox2* gene with predicted developmental regulatory elements (PADREs) across developmental stages. Colour codes as in **a**. **d**, Box plots showing the total number of PADREs per chromatin state across developmental stages indicated by dot shading. **e**, Stack bar of number of PADREs overlapping CAGE-defined promoters across six developmental stages and coloured as in **a. f**, Heatmap of pairwise Jensen-Shannon Divergence (JSD) of PADREs across developmental stages, **g,** Boxplots showing sequence conservation by phastCons scores (cons) of annotated PADREs.

Consensus peaks were called across all samples, thus each developmental stage shared the same set of 254,276 PADREs, with their chromatin annotation varying by stage (**Supplementary Table 12)**. The regulatory dynamics of PADREs are illustrated at the *sox2* locus, where chromatin state transitions across development were clearly visible (**Figure 3c**). The dominance of OpenChr and Quies categories indicates that only a fraction of all PADREs were active at any single developmental stage (**Figure 3d**). Quiescent PADREs — consensus peaks lacking both chromatin marks and accessibility signal at a given stage — represent elements not active at that stage. OpenChr PADREs, which were accessible but lacked the histone marks targeted in this study, may correspond to primed or decommissioned cis-regulatory elements or non canonical elements of unknown function. Similarly, PADREs annotated with flanking states (TssFlank1, TssFlank2, EnhFlank) carry histone modification signals but lack ATAC-seq support at that stage. In standard ChromHMM annotation, these states typically neighbour active regions; however, in the context of PADREs, they may represent poised, primed, or decommissioned elements.

Many promoter-annotated PADREs fell outside Ensembl annotated promoter-proximal regions, raising the possibility of mis-annotation (**Supplementary Figure 9**). Therefore we sought to validate these promoters by using Transcription Start Site (TSS) detection by CAGE-seq. We intersected PADREs with CAGE tag clusters and found that from 12,953 to 15,250 PADREs per developmental stage had TSS support (**Figure 3e, Supplementary Table 13**). The majority of CAGE-supported PADREs were annotated as TssA1 or TssA2; we designated these as high-confidence promoter PADREs, while promoter-annotated PADREs lacking CAGE support may represent enhancer mis-annotations. Although most high-confidence promoter PADREs overlap Ensembl TSS annotations, a subset did not fall in proximity to any annotated Ensembl TSS, representing potentially novel active promoters that could expand the current transcriptome annotations for the common carp (**Supplementary Figure 9**).

To assess relationships between regulatory landscapes across development, we computed Jensen–Shannon divergence (JSD) of PADRE activity, measured as ATAC-seq signal, across all stage pairs. This analysis revealed three distinct regulatory phases: an early phase encompassing late blastula and gastrula, a segmentation and early pharyngula phase, and a late pharyngula stage (**Figure 3f**).

Finally, we assessed the evolutionary conservation of PADREs using PhastCons scores derived from multispecies vertebrate alignments (**Figure 3g**). Polycomb-repressed regions and active enhancers exhibited the highest conservation, consistent with their known roles in conserved developmental gene regulation^23–26^. PADREs active in later developmental stages were more conserved than those active at earlier stages, mirroring observations in zebrafish and other vertebrates^14,27^, and consistent with the developmental hourglass model, which predicts that the highest developmental constraints occur at the so-called phylotypic stage towards mid-embryogenesis rather than at earlier stages such as blastula and gastrula^13^.

### Subgenome distribution of PADREs and its impact on gene regulation

To investigate the relationship between regulatory activity and subgenome origin, we examined the distribution of PADREs between the two carp subgenomes. We mapped the 254,276 PADREs on canonical chromosomes to homeologous coordinates (122,002 in subgenome A; 132,274 in subgenome B). We defined corresponding positions on homeologous chromosomes by pairwise whole-genome sequence alignment^28^ (“projections”), enabling direct comparison of regulatory potential between homeologous loci (**Figure 4a**). Based on this projection, we classified PADREs into three categories: homeologous PADREs, where the projected coordinates overlapped with PADREs in both subgenomes; asymmetric PADREs, where the PADRE can be projected but the alignment does not overlap a PADRE in the other subgenome; and singleton PADREs, which lack an alignable sequence in the homeologous locus (**Figure 4a, Supplementary Table 14**). Homeologous PADREs constituted the largest category, followed by singleton PADREs and asymmetric PADREs (**Figure 4a**). Notably, subgenome B carried a disproportionately higher number of asymmetric PADREs compared to subgenome A (**Figure 4b**), demonstrating regulatory asymmetry between the two subgenomes.

**Figure 4.**
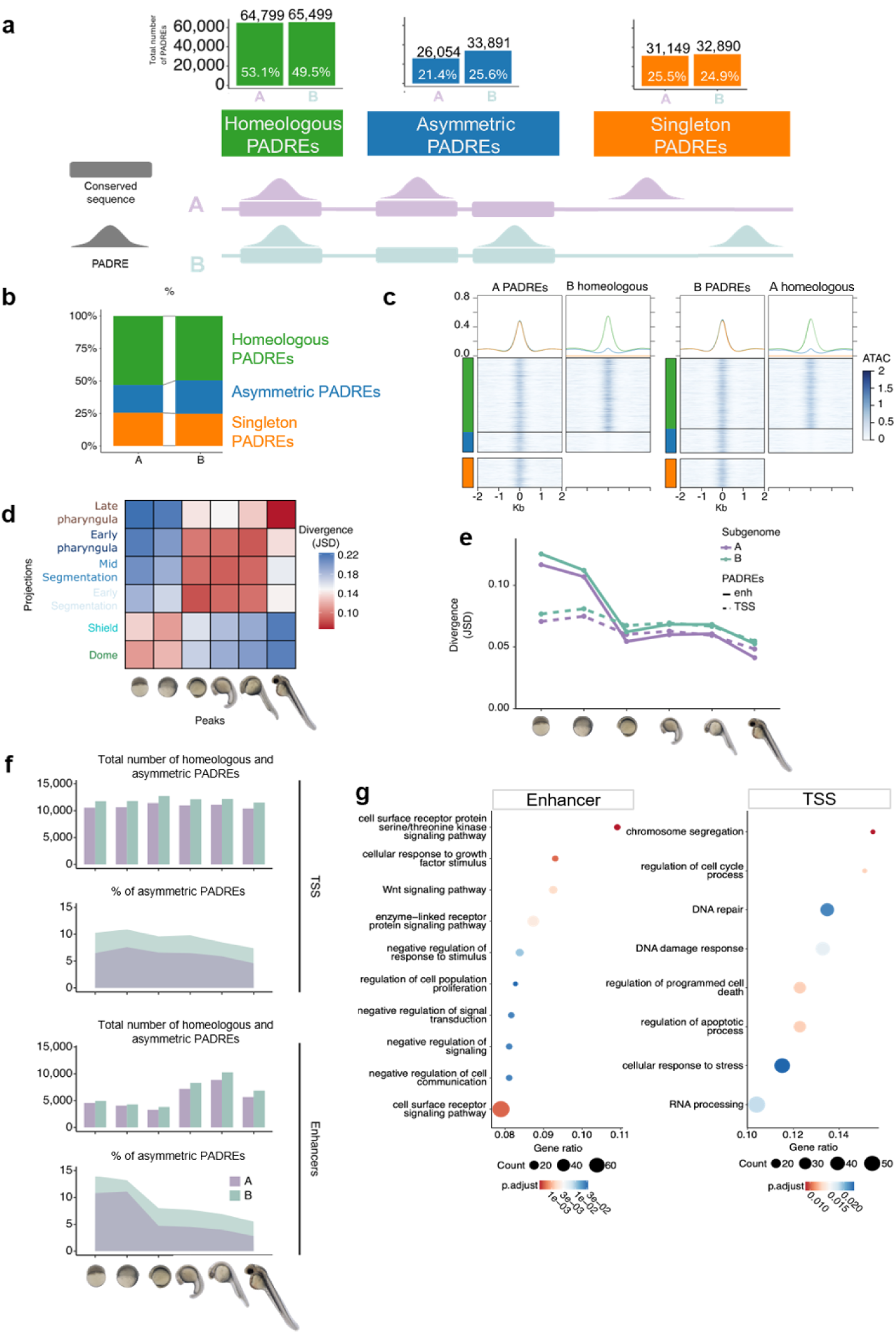
Subgenome-specific dynamics of chromatin accessibility across carp development. **a,** Schematic illustrating the categories of PADREs identified in homeologous regions. Numbers and percentages proportion of each category relative to the total are shown. **b**, Stack bar of percentage of PADREs classified in each homeology group, divided by subgenome. **c**, Heatmaps of ATAC-seq signal centered on homeologous PADREs for subgenomes A (left) and B (right) and their homeologous loci during mid-segmentation. Colors as in **a**. Aggregate signal profiles are on top. **d,** Heatmap of Jensen-Shannon distance (JSD) between TPMs of PADREs and their homeologs. **e**, Line plot of JSD over developmental time for subgenomes separating enhancer and TSS-associated PADREs. Values reflect the degree of divergence in chromatin accessibility between subgenomes. **f**, Bar charts show the total number of PADREs and regions homeologous to TSS. Area charts show the percentage of homeologous regions relative to the total number shown in the bar charts. **g**, Gene ontology (GO) plots of enriched biological process terms for differentially accessible PADREs between subgenomes.

To determine whether asymmetric PADREs reflect genuine subgenome-specific activity, we examined ATAC-seq signals at these loci and at their projections. These PADREs showed clear ATAC-seq signal in one subgenome but lacked signal at the corresponding homeologous coordinates (blue box in **Figure 4c**), confirming that these elements were uniquely active in one subgenome.

Then we asked the overall relationship between regulatory element variation between subgenomes and quantified regulatory divergence between homeologous elements at each developmental stage using Jensen–Shannon divergence (JSD) (**Figure 4d**). Divergence was highest at early stages and decreased progressively through development, with the early and late pharyngula stages showing the lowest JSD values. This developmental trend mirrored the convergence of transcriptome dynamics in **Figure 2c** and the three regulatory phases identified in **Figure 3e** and together are consistent with the hourglass model of embryonic development^12,13^. To determine whether this trend could be traced to either or both of the two main regulatory element types, we separately analysed promoter and enhancer PADREs (**Figure 4e**). Enhancers showed markedly higher divergence than promoters, particularly at early developmental stages, indicating that enhancer activity is less constrained between subgenomes than promoter activity.

Across all comparisons, subgenome B PADREs showed systematically higher divergence from their projections in subgenome A than in the reverse (**Figure 4e**), indicating asymmetric regulatory constraint.

Examination of PADRE numbers across development revealed distinct dynamics for promoters and enhancers (**Figure 4f, Supplementary figures 10,11**). Promoter PADREs were relatively stable in number across stages, with the proportion of subgenome-specific elements remaining constant and consistently higher in subgenome B. In contrast, enhancer PADREs increased substantially in number during development, while the proportion of subgenome-specific enhancers decreased progressively. The number of active enhancers in later stages was attributed to the activation of homeologous PADREs, since the % of asymmetric PADREs dropped drastically after early segmentation. Also, the trend was much more prominent in subgenome A, where in late pharyngula, only 2.5% of enhancers with homeology in subgenome B were active in subgenome A only. To quantify these dynamics, we fitted a binomial GLM modelling the proportion of subgenome-specific elements as a function of developmental stage, subgenome and element type, with element-level bootstrapping to validate significance (**Supplementary Table 15)**. The proportion of subgenome-specific elements declined during development (stage effect: β=-0.322, p<0.0001), and subgenome B elements declined significantly less steeply than A elements (stage × subgenome interaction: β= +0.110, 95%CI [0.075, 0.145], p<0.0001), indicating that subgenome A is more constrained. This differential constraint was significantly stronger for enhancers than for promoters (three-way interaction stage × subgenome × element type: β=−0.083, 95%CI[−0.126,−0.040], p<0.0001). The stage × subgenome interaction was four-fold larger for enhancers (+0.110) than for promoters (+0.027), demonstrating that the developmental convergence of subgenome activity is driven primarily by enhancers, while promoters maintain a stable baseline of B dominance with minimal developmental dynamics. An example for enhancer activity divergence at the *bmp7a* homeologous loci with corresponding gene expression variation between subgenomes **Figure 5a**.

**Figure 5.**
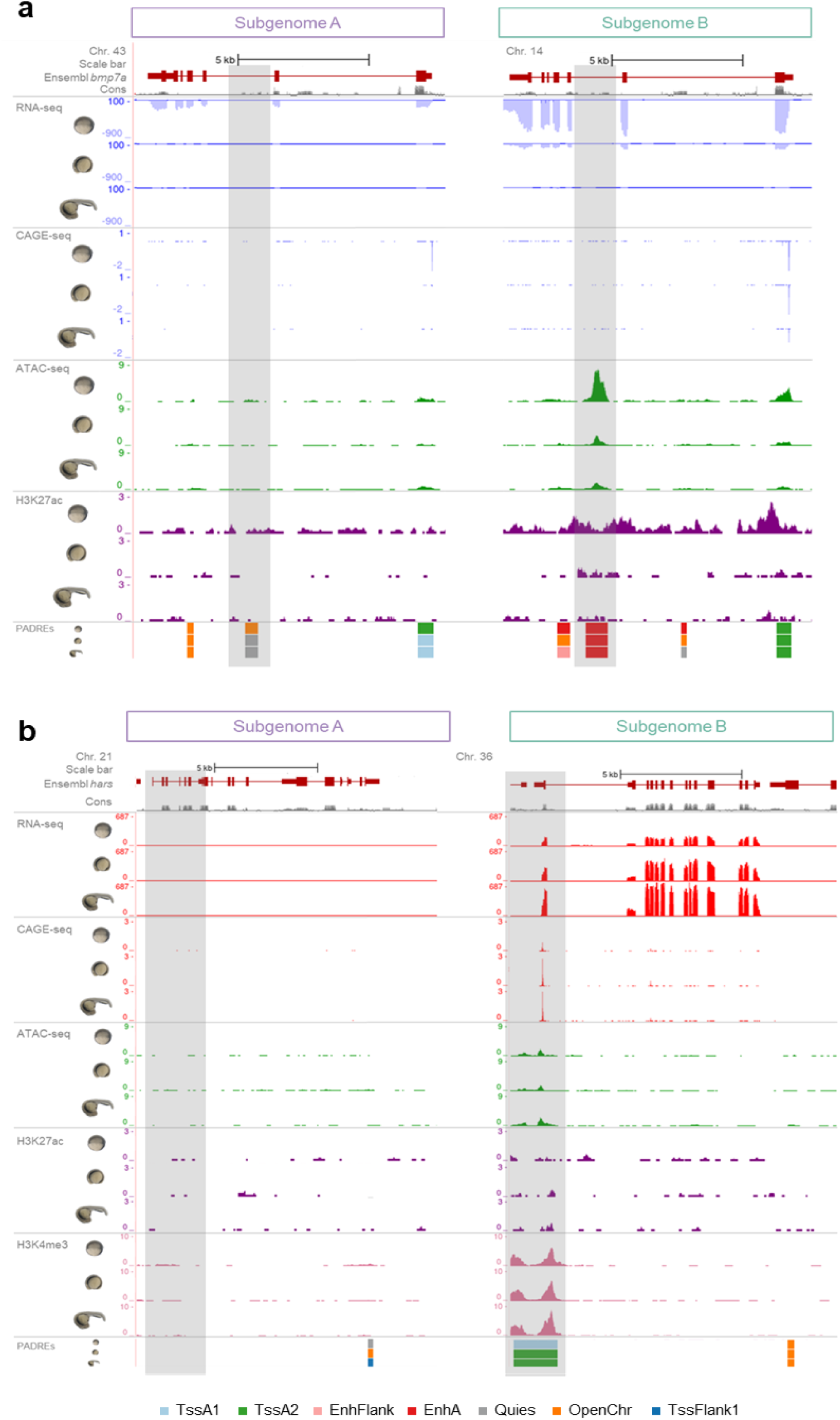
**a,** UCSC genome browser screenshot showing the developmental dynamics transcription (RNA-seq, CAGE-seq) and homeologous PADREs at *bmp7a* locus with chromatin feature support (ATAC-seq and H3K27ac). Grey bars highlight enhancer activity variation. **b,** UCSC genome browser screenshot showing the developmental dynamics of a PADRE in a homeologous region annotated as a TSS of the gene *hars*. CAGE-seq, ATAC-seq, H3K27ac and H3K4me3 tracks support the promoter annotation. Associated transcriptional activity (RNA-seq) shows gene expression differences between subgenomes. Grey bars highlight promoter feature variation.

Notably, promoters showed significant overall subgenome B dominance (subgenome × element type interaction: β = +0.277, p<0.001) but no developmental trend, ruling out a genome-wide technical artefact and confirming that the developmentally dynamic effect is specific to the enhancer regulatory layer. An example for developmentally constitutive promoter activity difference between subgenomes was seen at the *hars* homeolog loci (**Figure 5b)**.

The progressive decline in the proportion of subgenome-specific enhancers paralleled with the decreasing inter-subgenome JSD observed at the same developmental stages (**Figure 4d,e**), suggesting that subgenome regulatory convergence during the phylotypic period is primarily attributable to enhancer activity. The enhancer-dominant nature of varying degree of divergence during development is consistent with the notion of enhancers are more likely subjected to divergence - for example by subfunctionalization^29^ - than promoters^11^, whose turnover is more tightly coupled to gene retention and loss^30–32^.

Taken together, these results indicate that subgenome B was the dominant subgenome for retained regulatory activity: when a regulatory element was active in only one subgenome, it preferentially remained active in subgenome B. This dominance was more static at promoters but dynamically reinforced at enhancers as development progressed toward the phylotypic stage, again suggesting that the hourglass-like constraint acted on enhancers. Taken together, these observations support the notion that gene expression variation is reflected in subgenome-specific regulatory divergence, and illustrated by the examples of the *hars* and *bmp7a* homeologous gene loci (**Figure 5)**.

Gene Ontology enrichment analysis of genes linked to subgenome-biased PADREs revealed associations with functional categories including DNA repair and RNA processing, with partial overlap to the functional classes enriched in genes showing subgenome-biased expression (**Figure 4g**, **Figure 2d, Supplementary Tables 16, 17**).

### Cross-species conservation of annotated regulatory elements between carp and zebrafish

Our regulatory map of common carp offers an ideal platform for comparative genomic analysis of regulatory evolution among cyprinids. Given that carp and zebrafish diverged before the carp-specific whole-genome duplication, zebrafish provides both a phylogenetic outgroup for tracing regulatory divergence between carp subgenomes and an independently annotated reference for validating carp PADREs. To compare conservation of regulatory activity among cyprinids, we projected carp PADREs onto the zebrafish genome and assessed their overlap with zebrafish PADREs^14^, classifying them into four categories (**Figure 6a, Supplementary Tables 18, 19**): pan-orthologous PADREs, element present as PADREs in both carp subgenomes and zebrafish; sub-orthologous PADREs, which were present in one carp subgenome and zebrafish but absent from the other carp subgenome; carp-detected PADREs, which were alignable to zebrafish but not overlapping any zebrafish PADRE; and lineage-specific PADREs, which were not alignable to zebrafish.

**Figure 6.**
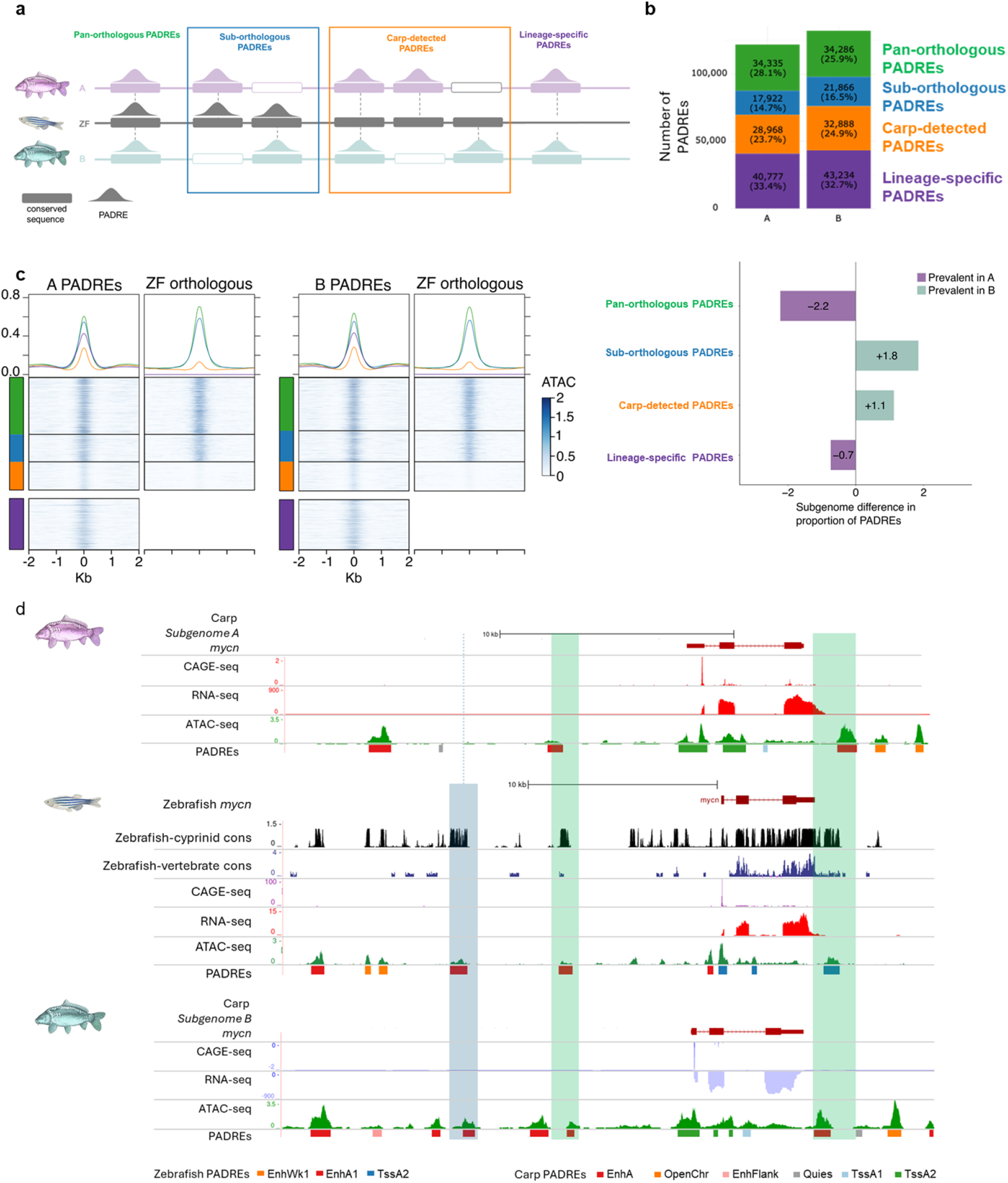
Functional conservation of regulatory elements with zebrafish. **a,** Schematic of PADRE categories identified through comparative analysis with zebrafish. **b,** Top, stacked bar chart showing the counts and proportions of the four PADRE categories for subgenomes A and B. Bottom, bar chart showing percentages of the subgenome difference in proportion of PADREs. **c**, Heatmaps of ATAC-seq signal centered at carp subgenome A PADREs (left pair) and subgenome B PADREs (right pair), showing carp subgenome signal alongside the corresponding zebrafish (ZF) signal. PADREs are grouped by category (coloured as in **a**). Aggregate signal profiles are above heatmaps. **d,** Genome browser screenshot of the *mycn* locus with annotated features and PADREs in zebrafish and carp subgenomes. Cons: Conservation tracks of zebrafish-cyprinid or zebrafish-vertebrate synteny. Green highlight: example of homeologous PADREs orthologous to zebrafish PADRE. Blue highlight: Example of subgenome B-specific PADRE, orthologous to zebrafish PADRE.

The distribution of PADREs across conservation categories differed significantly between subgenomes (chi-squared test, χ²=296.8, df=3, p<2.2×10⁻^16^), though effect sizes were modest (Cramér’s V=0.034), reflecting subtle but consistent compositional bias between subgenomes (**Figure 6b**). Subgenome A was proportionally enriched in pan-orthologus PADREs +2.2 percentage points (**Figure 6b**, bottom; proportion test, p<2.2×10⁻^16^), indicating that A’s regulatory repertoire is disproportionately composed of deeply cyprinid -conserved elements retained across both subgenomes. In contrast, subgenome B was enriched in sub-orthologous PADREs (+1.8pp, p<2.2×10⁻^16^) and carp-detected PADREs (+1.1pp, p=5.0×10⁻^11^). Lineage-specific PADREs were similarly represented in both subgenomes (∼33%). Together, these results suggest that subgenome B uniquely retained a greater proportion of ancestral regulatory elements than subgenome A, while subgenome A’s PADRE landscape has been stripped down to predominantly those elements conserved across both subgenomes. This asymmetric regulatory bias of subgenome A was consistent with the dominance of the B subgenome in common carp.

ATAC-seq signal profiling at projected loci further supported these classifications (**Figure 6c, Supplementary figure 12**). Pan-orthologous PADREs displayed the strongest ATAC-seq signal in both carp and zebrafish, while sub-orthologous PADREs showed a reduced but clear signal. Notably, carp-detected regions still showed weak but non-zero ATAC-seq signal in zebrafish and reduced signal in carp, indicating that many of these elements were likely functional, but fell below peak-calling thresholds, potentially due to low activity or cell-type specificity masked by bulk data. This implies that strict peak-overlap criteria underestimated the true extent of regulatory conservation, and that the fraction of functionally conserved elements may approach approximately two-thirds when weakly active orthologous regions were included.

A representative example is the *mycn* locus, where the subgenome B regulatory landscape was mirrored by the orthologous zebrafish locus (**Figure 6d** blue highlights). The conserved arrangement of PADREs and the concordant expression patterns of *mycn* in subgenome B and in zebrafish suggests that only subgenome B retained the cyprinid-ancestral regulatory architecture.

## Discussion

We present the first regulatory atlas of common carp embryogenesis, integrating transcriptomic and chromatin profiling across key developmental stages to expand existing annotations^20,33–35^. We provide a toolset for studying gene regulation in the most recent allotetraploid vertebrate. Public metadata, UCSC track hubs, and standardised pipelines make this a foundational reference for developmental and evolutionary studies in cyprinids. The resource also has aquaculture relevance: carp is among the most important freshwater aquaculture species globally but has lacked the functional regulatory annotation which will enable integration with mapping data to identify candidate variants underlying economically important traits^19^.

Common carp’s recent whole-genome duplication provides a unique opportunity to examine how subgenomes partitioned early regulatory programmes. Across 14,850 one-to-one homeolog pairs, subgenomes were transcriptionally coordinated through development, with asymmetry restricted to early stages, where subgenome B was more highly expressed — contrasting with prior reports of constitutive B dominance^20^ and consistent with the developmental hourglass model^15,27,36^. Chromatin segmentation shows that the core regulatory logic — active promoters and enhancers, polycomb repression, bivalency — are conserved with other vertebrates^14,37,38^ despite genome duplication, and the phylotypic stages showing the highest regulatory similarity^39^.

These transcriptomic and epigenomic patterns converge on a coherent model: subgenome-specific regulatory divergence peaks during early embryogenesis and contracts as both homeolog expression and PADRE states become increasingly coordinated toward the phylotypic stage. Previous work in *Xenopus laevis* showed that enhancer gain and loss drive subgenome asymmetry at zygotic genome activation^11^. To our knowledge, this is the first demonstration that the hourglass model extends to the subgenomes of an allotetraploid, with intragenomic regulatory divergence itself subjected to phylotypic constraint. Within this framework, the directional purging of enhancers in subgenome A — together with the higher zebrafish orthology of subgenome B PADREs — indicates progressive B dominance at the regulatory level, even when transcriptional output between homeologs becomes balanced.

Comparison with zebrafish shows that roughly two-thirds of carp PADREs are orthologous, reflecting strong conservation of cis-regulatory logic across cyprinid evolution despite carp-specific WGD, and consistent with deep conservation of organogenesis enhancers across vertebrates^40^. Lineage-specific PADREs likely reflect allotetraploidisation, subfunctionalisation, or divergence not captured by sequence-level orthology and warrant future studies to the investigation of global and gene specific mechanisms of regulatory divergence

Limitations of the study: Our histone-mark panel does not include H3K4me1 and other potentially useful marks for regulatory element classifications, and as a result, poised and primed enhancer states are likely underrepresented among open-chromatin regions without the H3K27ac mark. PADRE-to-gene assignment relies on proximity; chromatin conformation (Hi-C) or perturbation data would strengthen regulatory linkages. Distinguishing relaxed selection from true evolutionary distance between subgenomes, and relative to zebrafish, will require broader comparative analyses. Lack of signal for any of the chromatin features tested may have resulted from the masking effect of bulk data generation and cell type specific features may have been missed. Such lack of sensitivity can be mitigated by future development of single cell chromatin feature analyses tools for common carp.Functional validation of candidate elements will be essential to confirm developmental roles and clarify the mechanisms underlying subgenome-specific activity.

## Materials and Methods

### Animals

An inbred European common carp (Cyprinus carpio) strain originated from a cross between carp of Hungarian (R8 strain) and Polish origin (R3 strain)^41^ was used as breeder and for embryo collection in this project. On the day before sample collection, adult females from the R3xR8 strain were injected with pituitary extract to trigger ovulation. On the day of embryo collection sperm and eggs were obtained by abdominal massage and in vitro fertilisation (IVF) was performed at room temperature (RT) after sperm activation with water. Upon IVF, a pronase treatment was applied to the embryos following our Standard Operating Protocol (SOP) available in the FAANG repository (https://data.faang.org/api/fire_api/samples/UOB_SOP_Carp_embryo_collection_RNAseq_20210917.pdf,https://data.faang.org/api/fire_api/samples/UOB_SOP_Carp_embryo_collection_ATACseq_20210917.pdf,https://api.faang.org/files/protocols/samples/UOB_SOP_Carp_embryo_collection_ChIPseq_20260520.pdf) Briefly, the carp embryos were submerged in a 10 mg/ml pronase (Roche, cat no. 10165913103) solution at RT until chorions softened and started to break. The pronase solution was removed and embryos were washed in E3 medium. Dechorionated embryos were raised in agarose-coated petri dishes at 24°C.

### Embryo imaging and staging

Carp embryos were incubated at 24 C in E3 medium (5.0 mM NaCl, 0.17 mM KCl, 0.33 mM CaCl 2, and 0.33 mM MgSO 4 in 1 L sterilized distilled water), imaged by Leica M205 FA stereomicroscope, (Leica DFC 7000 T camera, Leica Application Suite X, Leica Microsystems GmbH; Wetzlar, Germany) under anaesthetic (100 mg/L MS-222 (tricaine methanesulfonate)). Stages of development were identified by detecting the embryonic features described for Danio rerio (PMID: 8589427) and naming of zebrafish stages were adapted to carp embryos exploiting the high morphological similarity between these cyprinid embryos. Timing of stages were detected as described in Supplementary Table 1.

### RNA sample preparation

Samples were collected in 700 µl of trizol and stored at −80° C. Total RNA was extracted using the miRNeasy mini kit (Qiagen, cat no. 217004) following the manufacturer’s instructions, also available as an SOP in the FAANG portal (https://api.faang.org/files/protocols/experiments/UOB_SOP_Carp_RNA_extraction_RNAseq_CAGEseq_20260506.pdf). Briefly, samples were lysed in RLT buffer passing the embryos through a needle and syringe. Lysates were passed through a MinElute spin column and centrifuged for 15 s at 13000g. Samples were washed with buffer RW1 and treated in-column with DNase I diluted in buffer RDD for 15 minutes at room temperature (RT). The digestion reaction was stopped with RW1 buffer and centrifuge step. Columns were washed with RPE buffer and 80% ethanol followed by a 5-minute centrifuge step to remove any ethanol residues. Finally, RNA was eluted in 14 μL of elution buffer. RNA integrity was confirmed on an RNA high sensitivity tape (Tapestation, Agilent), and concentration was determined with the RNA high sensitivity kit in qubit (invitrogen).

### RNA-seq library preparation

RNA library preparation was performed by Novogene UK. Poly-A selected RNA was obtained, and sequencing was carried out on an Illumina Novaseq 6000 (150 base pairs, paired-end reads). Library preparation and cDNA synthesis was performed using NEBNext Ultra Directional RNA Library Prep Kit for Illumina.

### ATAC-seq sample collection and nuclear preparation

Embryos at the desired stages were collected for nuclei preparation following the SOP available in the FAANG repository. Briefly, embryos were transferred to swelling buffer and dissociated by vigorous pipetting or pestle for early and late pharyngula stage embryos. The solution was filtered using a 50μm tube top filter and stored in a 15 ml tube. Samples were vortexed and kept on ice for 5 minutes. A centrifuge step was applied (500g for 5 minutes at 4°C), supernatant was removed and the remaining pellet was resuspended in 200μl of freezing buffer. At this step, samples can be stored for several months at −80°C or can be taken through to final steps of nuclear preparation. Continue the preparation by adding 1ml of cell lysis buffer, vortexing and keeping on ice for 5 minutes. Centrifuge the samples (500g for 5 minutes at 4°C) and remove all the supernatant. Resuspend the pellet in 500μl of cell lysis buffer and repeat the wash step spinning and removing the supernatant. Resuspend the pallet once more in 500μl of cell lysis buffer. At this point a QC step should be performed to confirm the integrity of the extracted nuclei. Transfer 10μl of the solution to a clean tube and mix with 5μl of Trypan blue. Confirm nuclear recovery efficiency with a cytometer.

### ATAC-seq library preparation

Chromatin accessibility was profiled using an adapted OmniATAC-seq protocol^42^, modified for carp embryo samples (https://api.faang.org/files/protocols/experiments/UOB_SOP_Carp_ATACseq_20260506.pdf). Nuclei isolated from carp embryos were resuspended in a transposase reaction mix comprising 2x TD buffer, transposase, PBS, and Tween-20. Tagmentation was carried out at 37°C for 30 minutes at 900 RPM, after which the reaction was stopped and DNA fragments were purified using a MinElute PCR purification kit and eluted in 21 µl EB buffer.

Transposed DNA was amplified using NEBNext Ultra II Q5 Master Mix with IDT Nextera UD indexes. An initial five-cycle pre-amplification was followed by qPCR-guided determination of additional cycle number, with amplification stopped at one quarter of maximum fluorescence to minimise PCR bias and GC/size artefacts. Final libraries were size-selected by double-sided AMPure XP bead purification to remove fragments smaller than ∼100 bp and larger than ∼670 bp, and eluted in TET buffer. Library concentration was quantified using a Qubit High Sensitivity assay, and fragment size distribution was validated by Tapestation (HS D5000 screentape), with successful libraries displaying a nucleosomal banding pattern with a predominant peak at approximately 200 bp.

### CAGE-seq library preparation

CAGE-seq followed a previously published protocol^43^ with sequencing compatibility optimisations. To improve sequencing efficiency and demultiplexing accuracy on Illumina platforms, the legacy 3-bp barcode at the start of Read 1 was replaced with 8-bp TruSeq Unique Dual Indexes. Adapter ligation and reverse transcription were followed by an additional five to six cycles of PCR amplification using NEBNext Ultra II Q5 Master Mix (New England BioLabs, #M0544S) to incorporate Illumina P5/P7 flow cell adaptors and I5/I7 indexes. Library purification was performed using 1. 4× AMPure XP beads (Beckman Coulter, #A63881) to remove adapter dimers and short fragments. Final libraries were quality-checked using a High Sensitivity DNA Bioanalyzer (Agilent, #5067-4626) and quantified with a Qubit dsDNA High Sensitivity Assay Kit (Thermo Fisher, #Q32854). Libraries including 512-cell, gastrula and early and mid segmentation stages were sequenced on an Illumina NextSeq 2000 with the flow cell P4-50 kit, generating 70 bp single-end reads. Libraries including the 2-cell stage and early and late pharyngula were sequenced on a NovaSeq 6000 v1.5 SP 100-cycle flow cell, generating 2 x 50 bp reads.

### ChIP-seq sample collection and library preparation

Samples collected in PBS with 1x cOmplete Protease inhibitor cocktail (Roche, cat. No. 04693116001). ChIP-seq libraries were prepared using the microchipmentation kit (Diagenode, cat no. C01011011) following the manufacturer’s instructions. An SOP is available in the FAANG repository(https://api.faang.org/files/protocols/experiments/UOB_SOP_Carp_ChIPseq_20260604.pdf). Input samples –non-precipitated chromatin – were produced in parallel and used as control for downstream analyses. The antibodies used for immunoprecipitation have been previously validated in zebrafish.

### NGS sequencing analysis

Genomics libraries (summaries in **Supplementary Tables 2-5**) have been analysed with standardised and freely available nf-core pipelines based on ENCODE^44^ guidelines. RNA sequencing samples were analysed using the nf-core RNA-seq pipeline^45^. This includes library QC (FastQC), sample trimming (Trim Galore!), read alignment (STAR) and quantification (RSEM). ATAC sequencing samples were analysed using the nf-core ATAC-seq pipeline (V 2.1.2). This includes library QC (FastQC), alignment to reference genome (BWA), intermediate QCs and peak calling (MACS2). ChIP sequencing samples were analysed using the nf-core ChIP-seq pipeline (V 2.0.0). This includes library QC (FastQC), alignment to reference genome (BWA), intermediate QCs and peak calling (MACS2). Cage sequencing analysis was carried out following sequencing; raw reads were demultiplexed using standard Illumina processing pipelines. Raw reads were aligned to the common carp reference genome assembly: Cypcar_WagV4. 0 (GCA_905221575. 1) using the nf-core cageseq 2. 0 pipeline (https://github.com/nf-core/cageseq/tree/dev, manuscript in preparation). Output tag clusters were further filtered to remove artefacts and low-expression peaks, and genomic coordinates were converted to a standardized format for quality validation of ChromHMM annotations.

### Genome-wide differential expression analysis

Raw count data were analysed using the DESeq2 package in R^46^. A DESeqDataSet object was constructed from the raw count matrix using the developmental stage as the design variable (design = ∼ Stage). Prior to analysis, low-count genes were removed by retaining only those genes with counts greater than 10. Normalisation and dispersion estimation were performed using the standard DESeq2 workflow. Pairwise differential expression comparisons were performed across consecutive developmental stages. Genes were considered significantly differentially expressed if they had an adjusted p-value ≤ 0.001 and a log₂ fold change ≥ 1.5.

### Homeolog gene pair differential expression analysis

To study subgenome bias in common carp we used a set of homeolog gene pairs (N = 14,849) with a 1:1 presence in subgenome A and B to compare the two subgenomes. To identify homeolog gene pairs, we obtained duplicated genes in common carp from Genomicus browser version 108. 01^47^ using the TreePattern Search tool with the following pattern: (Cyprinus. carpio. carpio:-1[<R>Cyprinus. carpio. carpio</R><S>1</S>],Cyprinus. carpio. carpio:-1[<R>Cyprinus. carpio. carpio</R><S>2</S>])D:-1;

We further filtered the list to include only 1:1 homeolog that are present on homologous chromosomes. The resulting homeolog set of 14,849 gene pairs contained strictly 1:1 homeolog shared between subgenome A and B. Homeolog set can be found in additional supplementary (**Supplementary Table 9**).

### Subgenome-specific differential expression analysis

To investigate transcriptional bias between the two subgenomes of the allotetraploid common carp genome, raw counts were first filtered to retain only the syntenic one-to-one homeolog pairs. The curated homeolog list was used to subset the full count matrix separately for subgenome A and subgenome B gene identifiers. These subsets were then merged with a gene ID-to-name reference table (derived from the genome annotation GTF file) to produce annotated count matrices for each subgenome. The two subgenome count matrices were subsequently joined on shared homeolog pair identifiers to create a single combined matrix containing count data for both homeologs of each gene pair.

Gene name harmonisation was performed to account for cases where only one homeolog carried a functional annotation. Specifically, where one homeolog lacked a gene name but its partner was annotated, the partner’s name was propagated across. Additionally, case-insensitive matching was applied to resolve naming inconsistencies between subgenomes. homeolog pairs in which both homeologs shared the same gene name were retained as high-confidence one-to-one pairs for downstream analysis; pairs with discordant names were recorded separately. Genes with zero counts across all samples were removed prior to DESeq2 analysis.

A DESeq object was constructed from the filtered homeolog count matrix, with each sample labelled to reflect both its developmental stage and subgenome of origin. Low-count filtering was applied as described above (counts > 10), and normalisation and dispersion estimation were performed using the standard DESeq2 workflow.

Within-stage subgenome bias was assessed by comparing subgenome A against subgenome B expression at each developmental stage using pairwise contrasts, covering all twelve stages from eggs through to pharyngula. Genes were considered to show subgenome bias if they met thresholds of adjusted p-value ≤ 0.001 and |log₂ fold change| ≥ 1, with positive fold changes indicating higher expression in subgenome A and negative fold changes indicating higher expression in subgenome B.

To identify genes with significant expression changes across all developmental stages simultaneously, a likelihood ratio test (LRT) was applied by fitting a full model (∼ Stage) against an intercept-only reduced model (∼ 1). Genes with an adjusted p-value < 0.01 were considered to vary significantly across development. Variance-stabilised counts (VST) for these genes were used to perform k-means clustering (k = 10, selected based on an elbow plot approach) via pheatmap^48^, grouping genes with similar expression trajectories across subgenomes and developmental stages. Principal component analysis was performed on the full VST-transformed count matrix to visualise overall sample relationships.

### Differential chromatin accessibility analysis

Chromatin accessibility between projected regions and their corresponding observed ATAC-seq peaks was assessed using edgeR (v4.4.2)^49^. The analysis was performed separately for active promoters (TssA1 and TssA2 chromatin states) and active enhancers (EnhA chromatin state) at each of six developmental stages. Read counts for both observed consensus peaks and their projected regions into the other subgenome were obtained with featureCounts and combined into a single count matrix per stage, with replicate identity included as a blocking factor in the design matrix (∼ replicate + class), where class levels are either peak or projection. Elements with a count lower than 5 were filtered out, libraries were normalized, and dispersion was estimated. Differential accessibility was tested using a likelihood ratio test, and elements were considered significantly differentially accessible at FDR ≤ 0.05 and |log₂FC| ≥ 2. The results were additionally stratified by subgenome (A or B) and by homeology class (homeologous PADREs or homeologous regions) to evaluate accessibility differences between subgenomes.

### Identification of promoters

Active promoters were identified by intersecting the consensus ATAC-seq peak set with ChromHMM chromatin state annotations. Peaks overlapping the active transcription start site states TssA1 and TssA2 were retained as promoter-associated differentially accessible regulatory elements at each of the six developmental stages. Promoter PADREs were further validated by overlap with CAGE-derived transcription start sites, providing independent support for active promoter activity. Subgenome assignment of each peak was performed based on chromosomal coordinates using the A and B subgenome chromosome maps of C. carpio.

### Identification of PADREs

Differentially accessible regulatory elements (PADREs) were defined as consensus ATAC-seq peaks annotated to active chromatin states by ChromHMM. The ChromHMM model was trained on the six developmental stages using matched ATAC-seq and ChIP-seq signal tracks, and a dense segmentation was produced for each stage. Peaks were assigned the chromatin state with the highest priority among overlapping states, following a defined priority order. These states were annotated following the Roadmap Epigenomics Project framework and Baranasic et al. (2022) ^14,50^ and defined as follows: TssA1, TssFlank1, TssA2, TssFlank2, EnhA, EnhFlank, OpenChr, Bival, PrC, Quies. A total of 255,028 PADREs were identified across the genome, of which 254,276 were located on canonical chromosomes and 752 on non-canonical sequences.

Only PADREs on canonical chromosomes were retained for subsequent homeology analyses. To identify relationships between PADREs across the two subgenomes, PADREs from subgenome A were projected onto subgenome B coordinates using a liftOver chain constructed from the pairwise alignment of the two subgenomes, and vice versa. Projections were reduced to single-interval representations (merging intervals within 300 bp) and filtered to retain only those mapping to the expected homeologous chromosome, ensuring that comparisons were made exclusively between syntenic regions. Based on the projection outcome, PADREs were classified into three groups: homeologous PADREs (present and overlapping as a PADRE in both subgenomes), asymmetric PADREs (present as a PADRE in one subgenome and projectable to the other, but not a PADRE at the projected location), and singleton PADREs (present in one subgenome but not projectable to the other).

### Zebrafish comparison

To assess cross-species conservation of carp regulatory elements, PADREs from each subgenome were projected onto the zebrafish (Danio rerio) genome using a stitched liftOver chain derived from whole-genome alignment of C. carpio(Cypcar_WagV4.0, GCA_905221575.1) to the zebrafish reference. Projections were performed separately for subgenome A and B PADREs, retaining only projections that mapped to a single contiguous interval and filtered to retain only those mapping to the orthologous zebrafish chromosome. Projected carp PADREs were then intersected with a set of consensus zebrafish PADREs (cPADREs). Each carp PADRE was classified into one of four categories based on the combined outcome of its carp-to-carp and carp-to-zebrafish projections: (i) homeologous PADREs orthologous to ZF PADREs — present as a homeologous PADRE pair in both carp subgenomes and overlapping a zebrafish cPADRE; (ii) subgenome-specific PADREs orthologous to ZF PADREs — present in only one carp subgenome but overlapping a zebrafish cPADRE; (iii) PADREs aligned to ZF — projected to the zebrafish genome but not overlapping a zebrafish cPADRE; and (iv) homeologous PADREs non-alignable to ZF — present as a homeologous PADRE pair in both carp subgenomes but not projectable to the zebrafish genome. The distribution of categories between subgenomes A and B was compared using a chi-squared test of independence with Cramér’s V as an effect size measure, and per-category differences in subgenome proportions were assessed with two-proportion z-tests with Bonferroni correction.

## Author contributions

AJG Designed experiments, carried out all embryo and genomics work and analysed genomics datasets, AMP analysed CAGE-seq datasets and performed PADREs analyses, YH generated CAGE libraries, and contributed to genomics data analysis, BZ generated CAGE libraries, ZCB and TM staged embryos and produced embryo images, AB, HJM and GW generated carp genome assembly, BL advised on genomics data analysis, DB designed and carried out genomics data analyses and genome segmentation, FM conceived the study. AJG, AMP, DB, and FM wrote the ms and all authors commented and approved the ms.

## Acknowledgements

This work was funded by the European Commission’s Horizon 2020 programme through the AQUA-FAANG (grant agreement no. 817923; to FM and BL) and PrecisionTox (grant agreement no. 965406; to FM) projects. TM was supported by the National Research, Development and Innovation Office of Hungary (NKFI ADVANCED 150916) and by the Flagship Research Group Programme of the Hungarian University of Agriculture and Life Sciences. This research was supported by the ‘PFAQuatic’ project (2024-1.2.3-HU-RIZONT-2024-00100), implemented with support from the Ministry of Culture and Innovation of Hungary through the National Research, Development and Innovation Office, financed under the HU-rizon funding scheme. ZC-B was supported by the Research Excellence Programme of the Hungarian University of Agriculture and Life Sciences. BZ was funded by the European Union – NextGenerationEU (grant NPOO.C3.2.R2-I1.06.0024) and the Croatian Science Foundation (project IP-2024-05-5224). DB and AMP were supported by the Horizon Europe MSCA programme through DANIO-ReCODE (grant agreement no. 101169349). DB was further supported by the European Union – NextGenerationEU through the CroAGE grant (NPOO.C3.2.R2-I1.06.0060) and by EMBO through an Installation Grant (EMBO-IG-6018). We thank Bence Ivanovics for assistance with preparing the figures.

## Data accessibility

All sequencing datasets produced in this study are available through the FAANG data portal (https://data.faang.org/projects/AQUA-FAANG) and the European Nucleotide Archive (ENA). The ENA accession numbers for the different studies are as follows: RNA-seq PRJEB112855, ATAC-seq PRJEB114341, ChIP-seq PRJEB114353 and CAGE-seq PRJEB115120. The code developed for the analyses and figures presented in this manuscript can be found at: https://github.com/mlkr-rbi/carp-regulatory-atlas.

## Notes

### Competing Interest Statement

The authors have declared no competing interest.

