## Supplementary Figure for "Mapping non-coding functional elements in allotetraploid *Cyprinus carpio* embryo development reveals subgenome variation of transcription regulation"

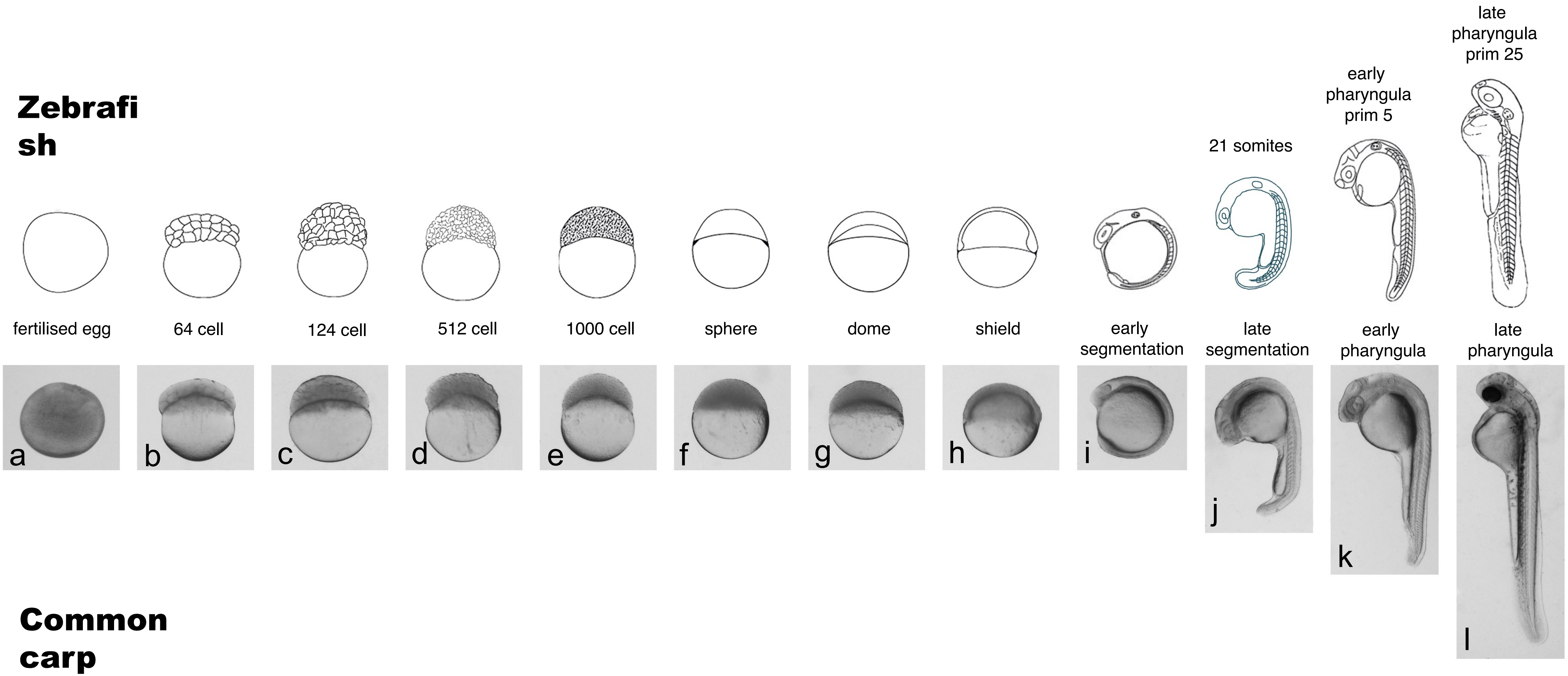


**Supplementary figure 1. Stages of early embryonic and larval development in zebrafish and carp.** Schematic diagrams (top) and corresponding stereoscope images (a–l, bottom) illustrate sequential developmental stages from fertilization through late pharyngula. (a) Fertilised egg; (b) 64-cell stage; (c) 124-cell stage; (d) 512-cell stage; (e) 1000-cell stage; (f) sphere stage; (g) dome stage; (h) shield stage; (i) early segmentation; (j) late segmentation (21 somites); (k) early pharyngula (prim-5); (l) late pharyngula (prim-25). Developmental stage names follow standard zebrafish staging nomenclature.

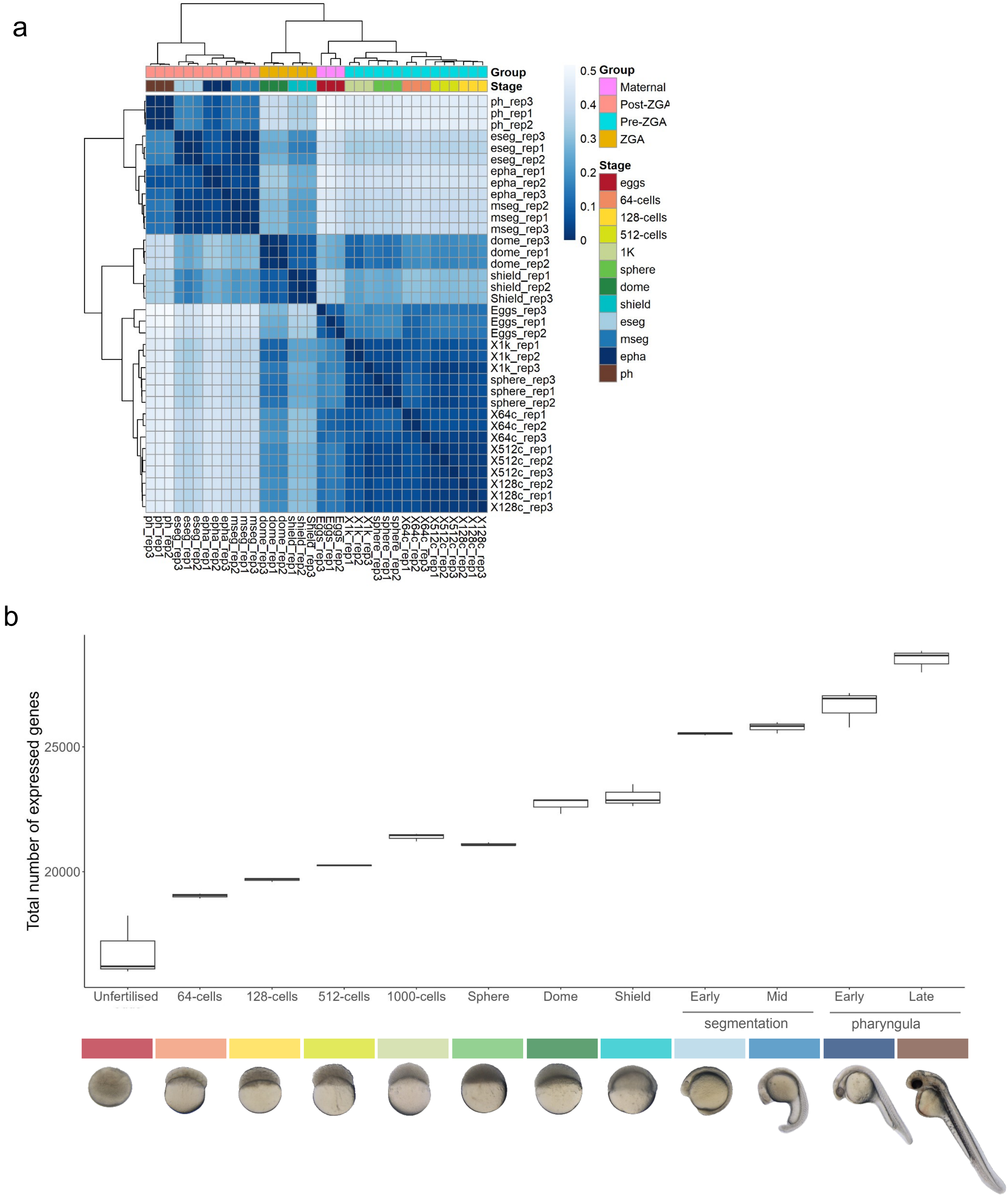


**Supplementary figure 2. a,** Sample correlation matrix of Spearman correlation coefficients of RNA-seq samples compared to each other. **b,** Box plots showing the total number of expressed genes at successive developmental stages, from unfertilised eggs through to the late pharyngula stage. n = 3 biological replicates per stage.

a

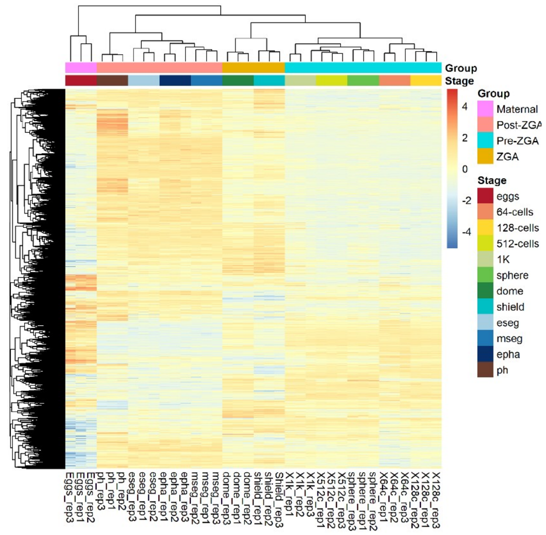


b

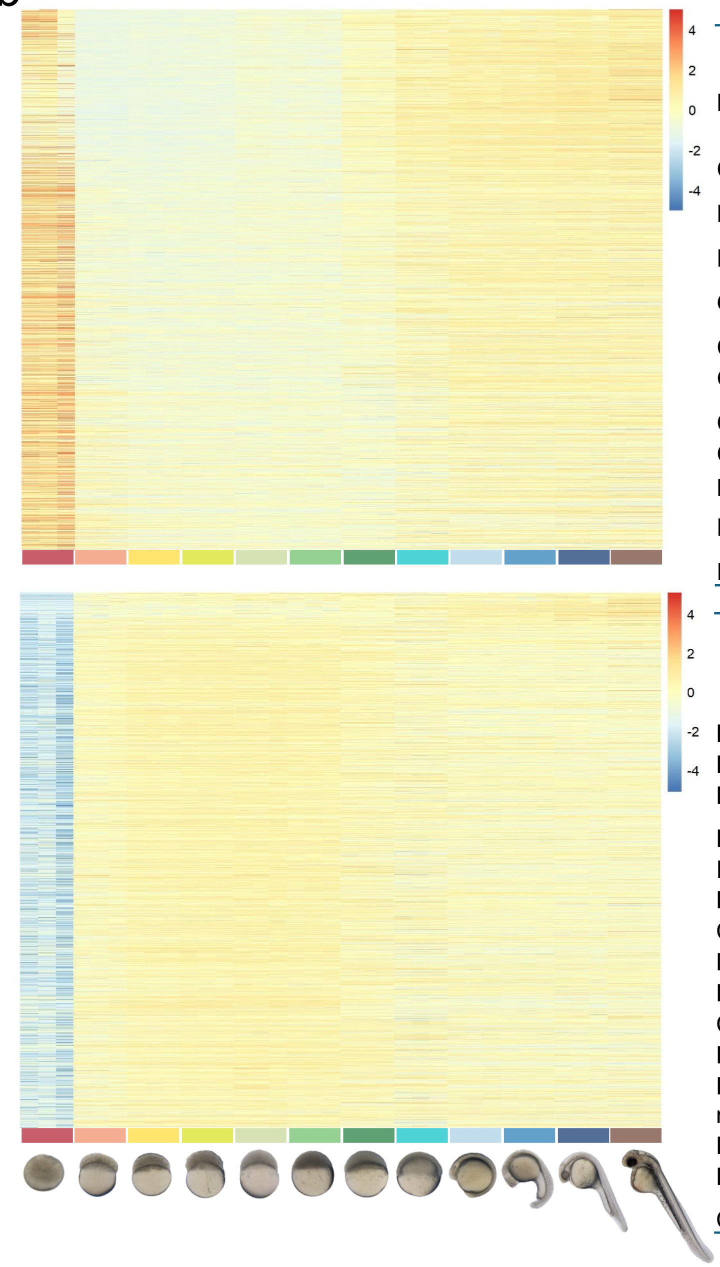

**Unfertilised egg upregulated genes (Egg/64-cells)**

| **Pathway** | **Enrichment FDR** |
|---|---|
| Cellular macromolecule metabolic proc. | 0.0007 |
| Biosynthetic proc. | 5.48E-10 |
| Protein metabolic proc. | 5.23E-06 |
| Organic substance biosynthetic proc. | 7.96E-10 |
| Cellular biosynthetic proc. | 1.44E-10 |
| Gene expression | 8.37E-07 |
| Cellular protein metabolic proc. | 5.01E-09 |
| Cellular nitrogen compound biosynthetic proc. | 2.38E-10 |
| Macromolecule biosynthetic proc. | 7.964E-10 |
| RNA metabolic proc. | 0.009E59529 |

**64-cell stage upregulated genes (64-cells/Eggs)**

| **Pathway** | **Enrichment FDR** |
|---|---|
| Protein modification by small protein | 2.77E-06 |
| Protein transport | 2.95E-05 |
| Protein localization | 1.48E-05 |
| Establishment of protein localization | 4.02E-05 |
| Intracellular transport | 2.95E-05 |
| Protein modification proc. | 4.69E-17 |
| Cellular protein modification proc. | 4.69E-17 |
| Macromolecule modification | 4.68E-17 |
| Protein phosphorylation | 2.12E-07 |
| Cellular protein metabolic proc. | 6.08E-18 |
| Phosphorylation | 4.12E-07 |
| Phosphate-containing compound metabolic proc. | 8.47E-09 |
| Phosphorus metabolic proc. | 1.02E-08 |
| Protein metabolic proc. | 3.87E-11 |
| Cellular macromolecule metabolic proc. | 3.85E-09 |

**Supplementary figure 3. Gene expression analysis across carp embryonic developmental stages. a.** Heatmap of gene expression across developmental stages following a likelihood ratio test analysis. **b.** Heatmap of upregulated genes in unfertilised eggs (top) and upregulated in 64-cell stage embryos compared to eggs (bottom, representing the first zygotic transcripts) and their GO biological process-related terms..

a

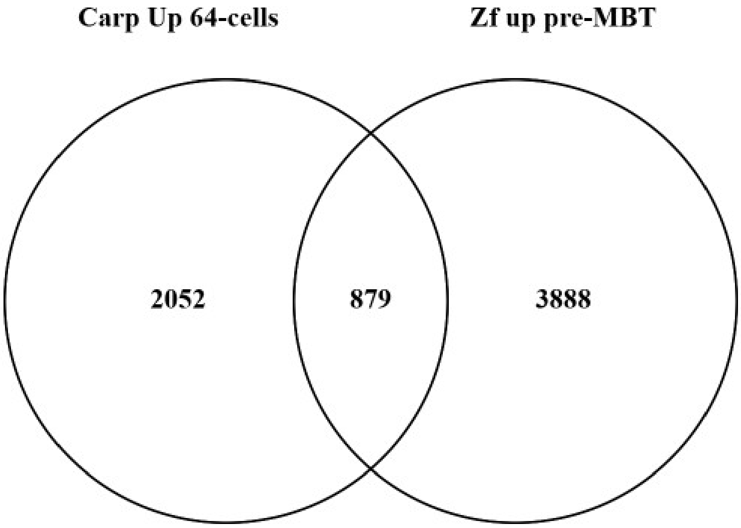


b

**Carp 64-cell stage upregulated genes (64-cells/Eggs)**
GO:0070647 protein modification by small protein
GO:0015031 protein transport
GO:0008104 protein localization
GO:0045184 establishment of protein localization
GO:0046907 intracellular transport
GO:0036211 protein modification proc.
GO:0006464 cellular protein modification proc.
GO:0043412 macromolecule modification
GO:0006468 protein phosphorylation
GO:0044267 cellular protein metabolic proc.
GO:0016310 phosphorylation
GO:0006796 phosphate-containing compound metabolic
GO:0006793 phosphorus metabolic proc.
GO:0019538 protein metabolic proc.
GO:0044260 cellular macromolecule metabolic proc.

**Zebrafish pre-MBT upregulated genes**
GO:0042254 ribosome biogenesis
GO:0022613 ribonucleoprotein complex biogenesis
GO:0006886 intracellular protein transport
GO:0015031 protein transport
GO:0045184 establishment of protein localization
GO:0044265 cellular macromolecule catabolic proc.
GO:0006396 RNA processing
GO:0009057 macromolecule catabolic proc.
GO:0046907 intracellular transport
GO:0071705 nitrogen compound transport
GO:0008104 protein localization
GO:0044248 cellular catabolic proc.
GO:1901575 organic substance catabolic proc.
GO:0009056 catabolic proc.
GO:0006996 organelle organization
GO:0071840 cellular component organization or biogenesis
GO:0044260 cellular macromolecule metabolic proc.
GO:0046483 heterocycle metabolic proc.
GO:0006139 nucleobase-containing compound metabolic proc.
GO:0006725 cellular aromatic compound metabolic proc.

**Supplementary figure 4. Comparison of upregulated genes in carp 64-cell stage and zebrafish pre-MBT stage embryos.** a, Venn diagram showing the overlap between genes upregulated in carp at the 64-cell stage relative to eggs and genes upregulated in zebrafish prior to the mid-blastula transition. B, Gene Ontology (GO) terms associated with the carp 64-cell stage upregulated gene set (left) and the zebrafish pre-MBT upregulated gene set (right), listed with corresponding GO identifiers.

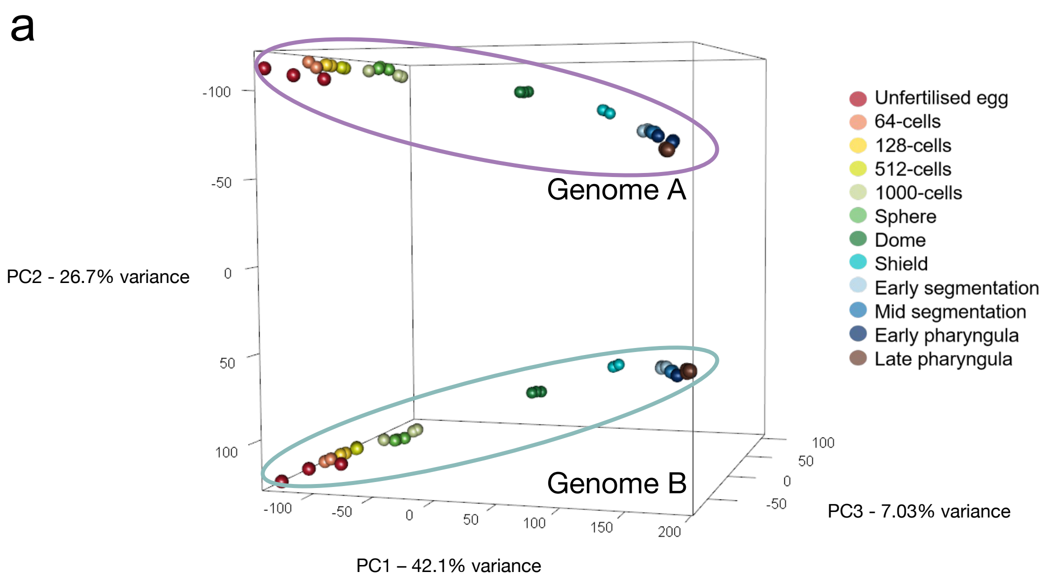


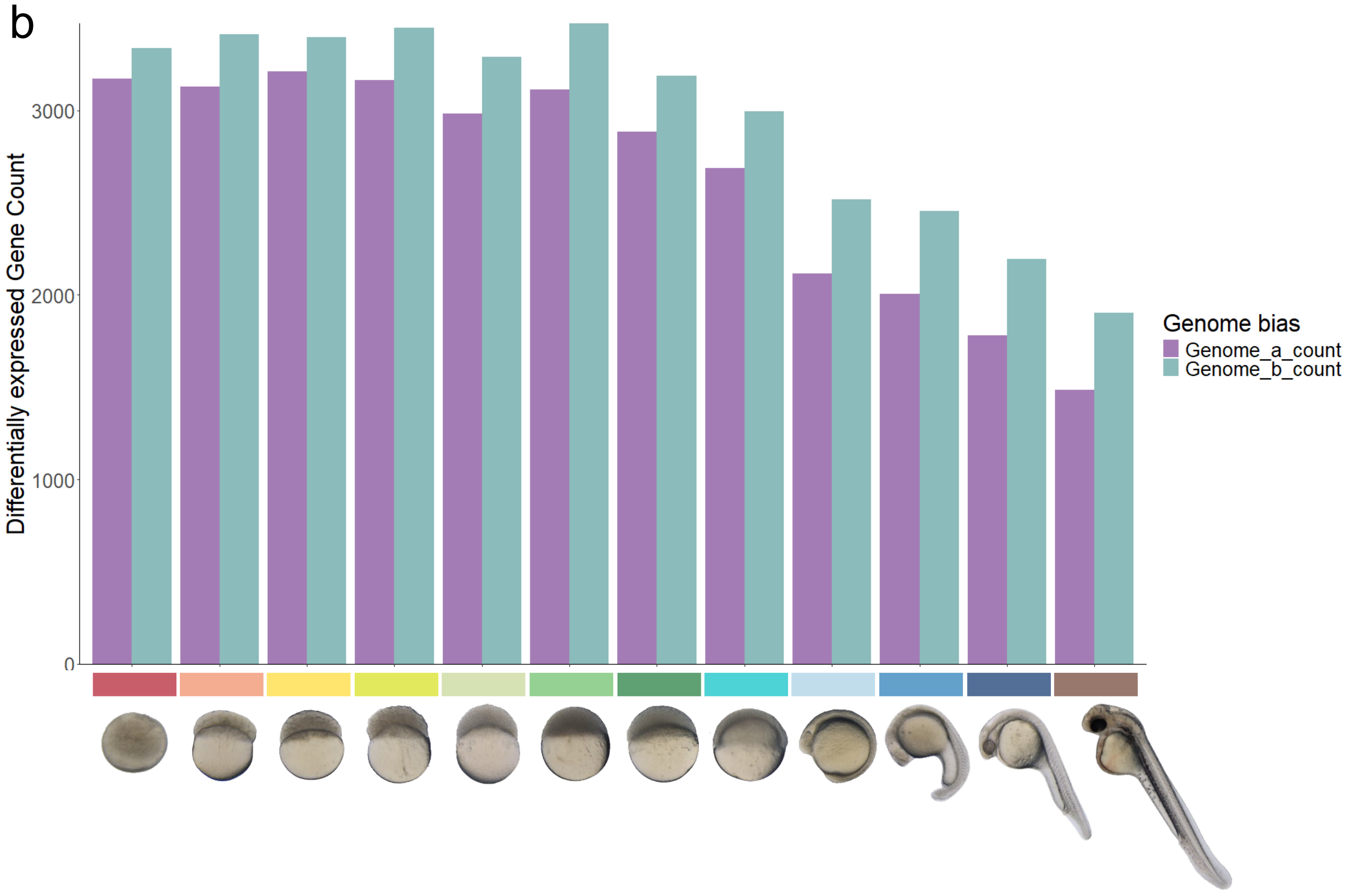


**Supplementary figure 5. Transcriptomic analysis of carp embryonic development across the two subgenomes. a,** Principal Component Analysis (PCA) of gene expression profiles across 12 developmental stages, from unfertilised egg to prehatch. Each point represents an individual biological replicate, coloured by developmental stage. Ellipses denote the two subgenome clusters (Genome A and Genome B). **b,** Bar chart showing the number of differentially expressed genes (DEGs) assigned to each subgenome across developmental stage comparisons. Each pair of bars corresponds to a developmental stage.

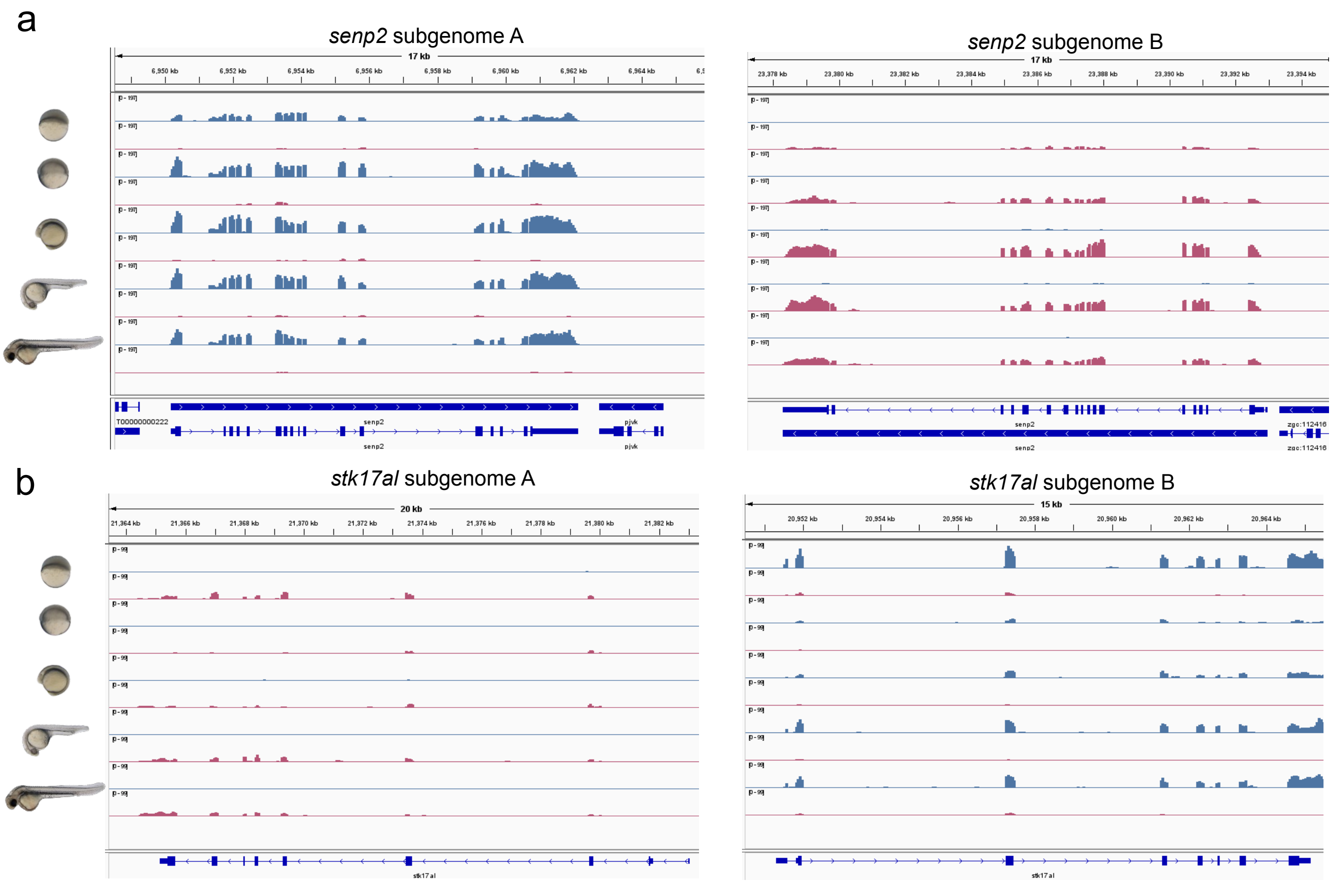


**Supplementary figure 6. Subgenome-specific expression of homeolog genes during carp embryonic development.**
**a.** IGV tracks showing RNA-seq read coverage at the *senp2* locus for subgenome A (left) and subgenome B (right). Each track corresponds to a developmental stage. The gene model is displayed at the bottom of each panel. **b.** IGV tracks showing RNA-seq read coverage at the *stk17al* locus for subgenome A (left) and subgenome B (right), following the same plotting conventions as in (b). Gene models are shown at the bottom of each panel.

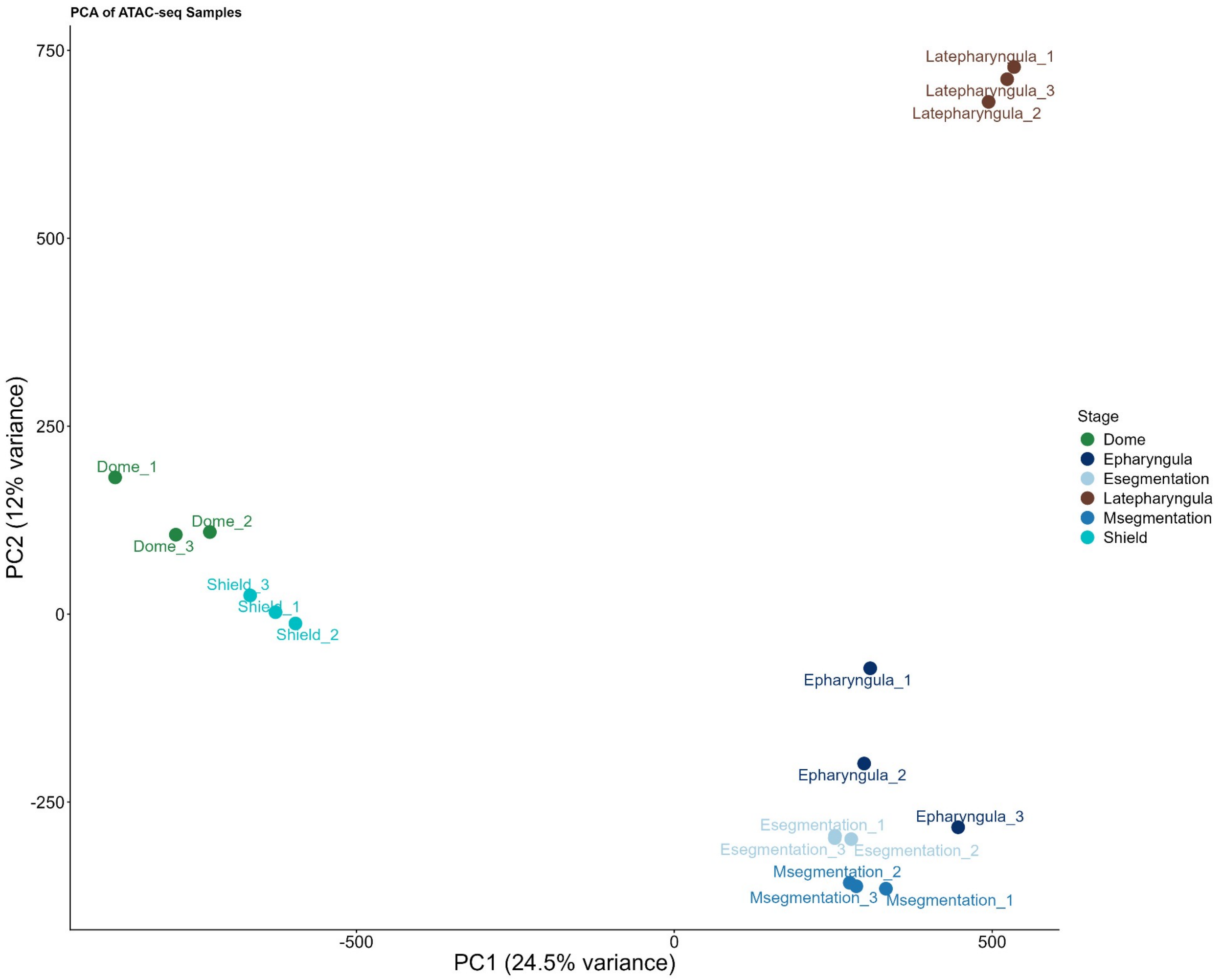


**Supplementary figure 7. Principal component analysis (PCA) of ATAC-seq samples across carp developmental stages.** Each point represents an individual replicate, coloured by developmental stage

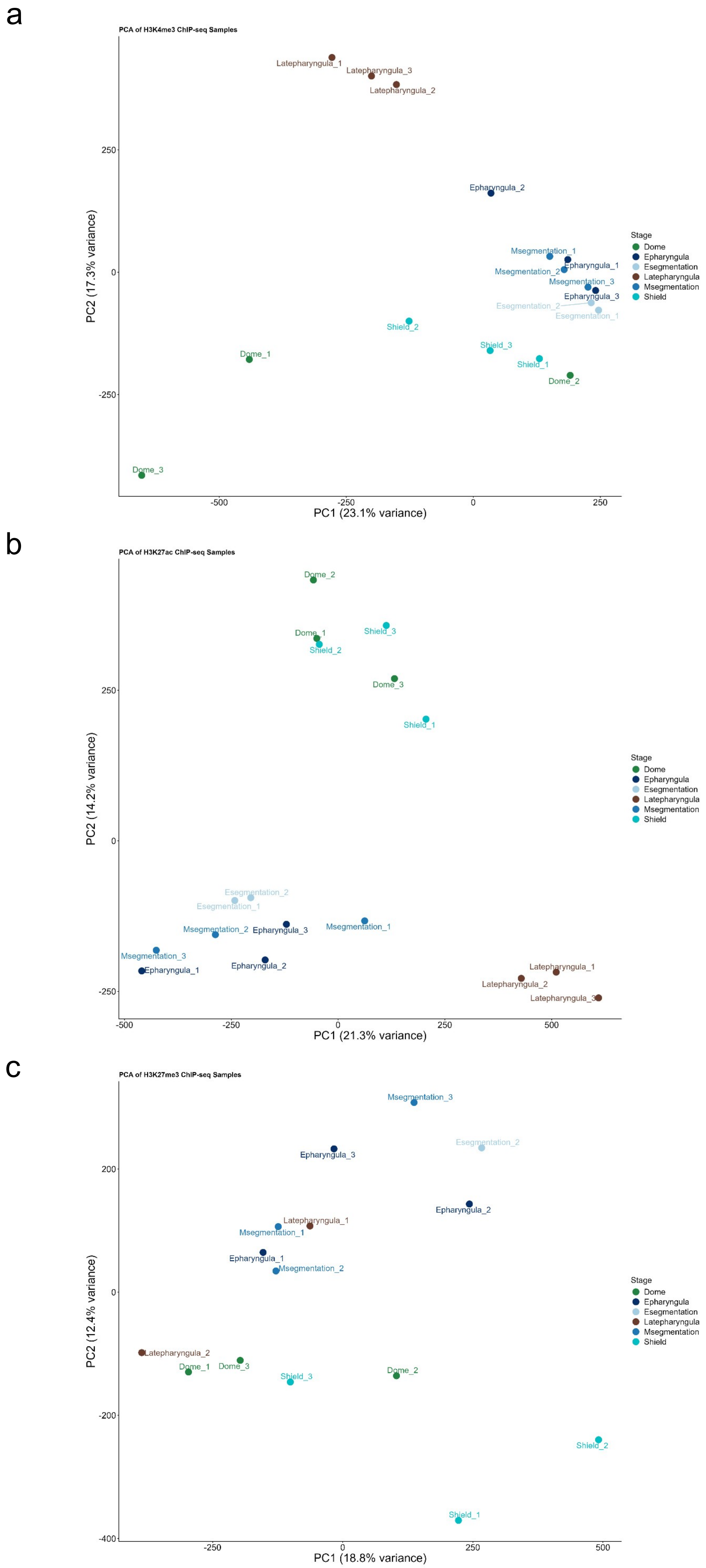


**Supplementary figure 8. Principal component analysis (PCA) of ChIP-seq samples for H3K4me3 and H3K27ac across carp developmental stages.** a, PCA of H3K4me3 samples. b, PCA of H3K27ac samples. c, PCA of H3K27me3 samples  Samples are coloured by developmental stage.

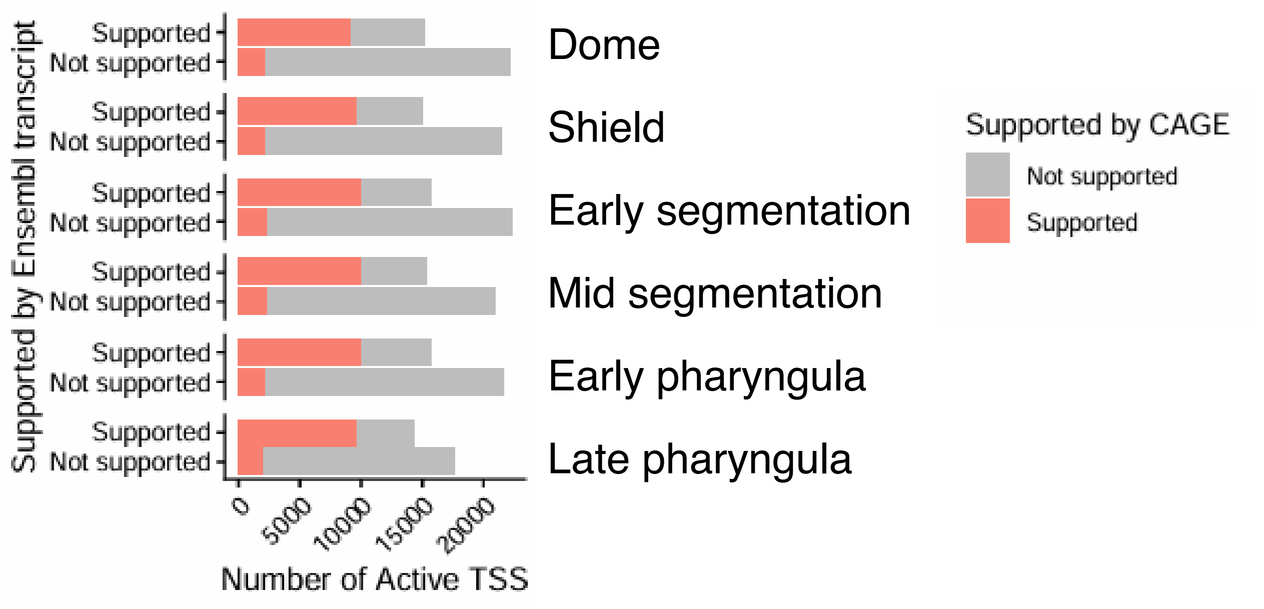


**Supplementary Figure 9. Validation of transcription start sites (TSS) by CAGE across carp developmental stages.** For each stage, active TSS are grouped by whether they are supported (top bar) or not supported (bottom bar) by Ensembl transcript annotation, and further classified by CAGE support (red, supported; grey, not supported). The number of active TSS is shown on the x-axis.

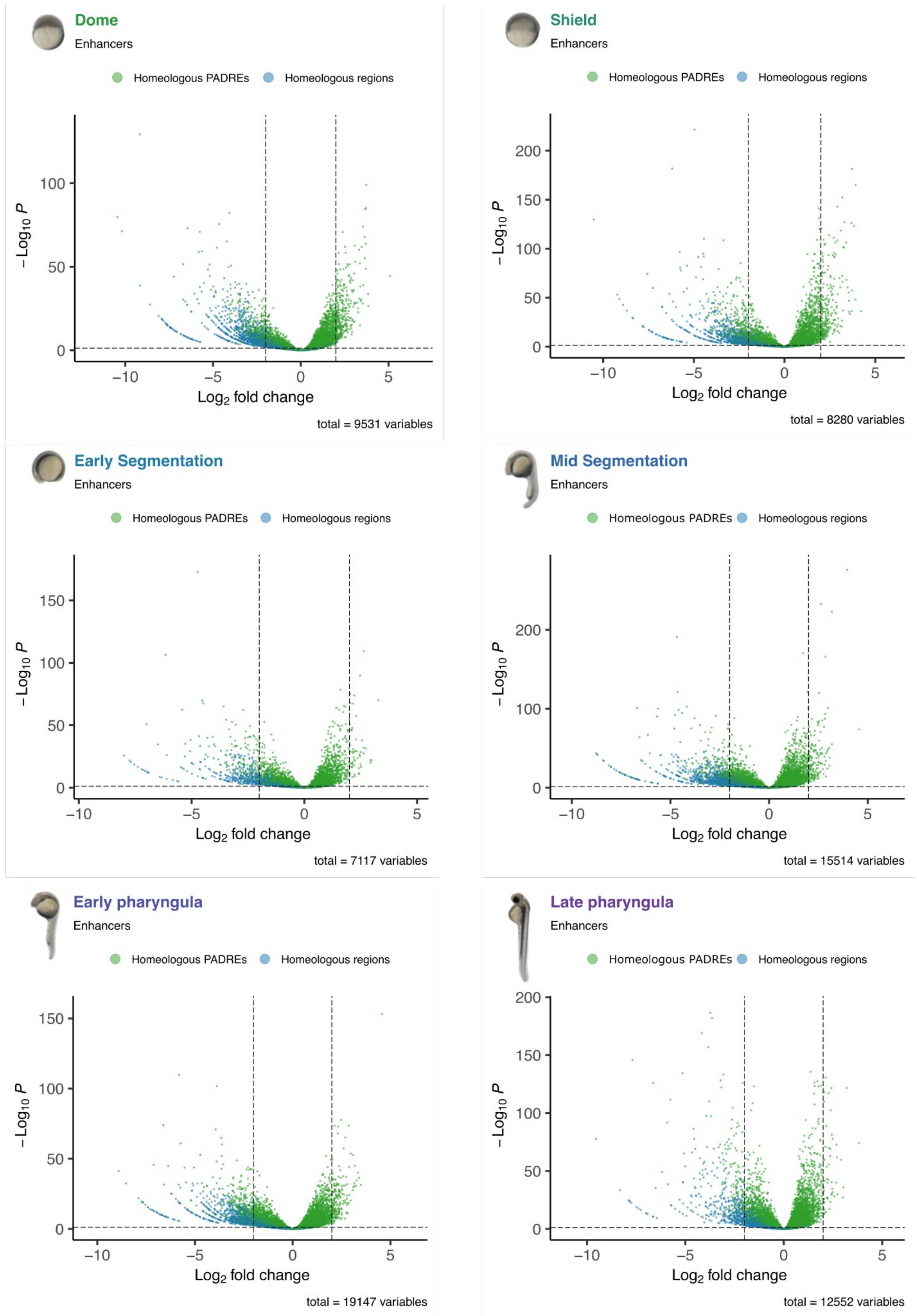


**Supplementary figure 10. Differential chromatin accessibility at homeologous enhancer PADREs across developmental stages.** Volcano plots showing differential chromatin accessibility between observed ATAC-seq peaks and their projected coordinates in the reciprocal subgenome, for enhancer PADREs at six developmental stages: Dome, Shield, Early Segmentation, Mid Segmentation, Early Pharyngula, and Late Pharyngula. The x-axis shows the $\log_2$ fold change and the y-axis the $-\log_{10}$ p-value. Each point represents a PADRE, coloured by homeology class: homeologous PADREs (green) and homeologous regions (blue). Dashed vertical lines indicate the $\log_2$ fold change threshold ($|\log_2 FC| \geq 2$) and the dashed horizontal line indicates the significance threshold ($FDR \leq 0.05$). The total number of elements tested at each stage is indicated below each plot.

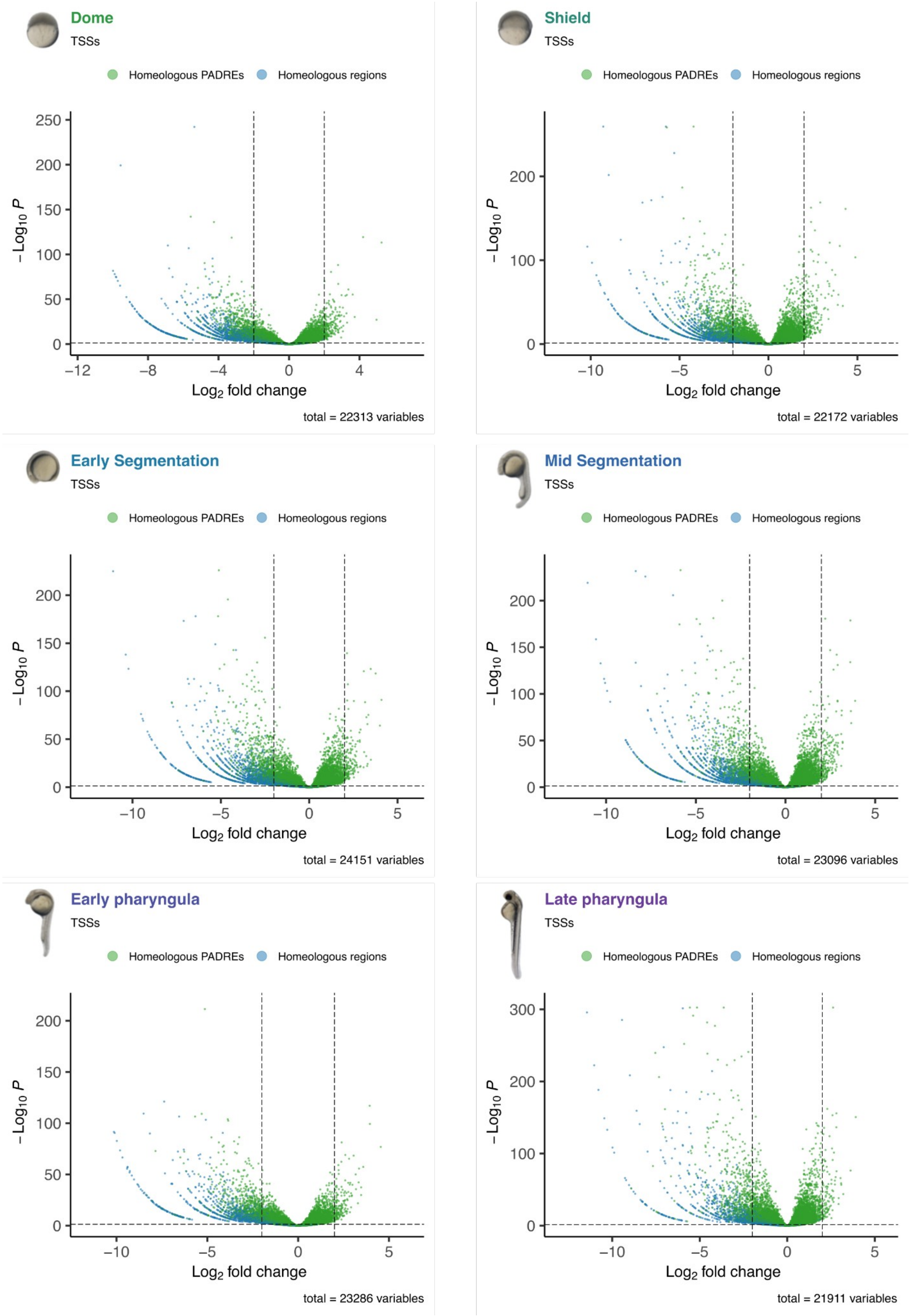


**Supplementary figure 11. Differential chromatin accessibility at homeologous TSSs PADREs across developmental stages.** Volcano plots showing differential chromatin accessibility between observed ATAC-seq peaks and their projected coordinates in the reciprocal subgenome, for TSS PADREs at six developmental stages: Dome, Shield, Early Segmentation, Mid Segmentation, Early Pharyngula, and Late Pharyngula. The x-axis shows the $\log_2$ fold change and the y-axis the $-\log_{10}$ p-value. Each point represents a PADRE, coloured by homeology class: homeologous PADREs (green) and homeologous regions (blue). Dashed vertical lines indicate the $\log_2$ fold change threshold ($|\log_2 FC| \geq 2$) and the dashed horizontal line indicates the significance threshold (FDR $\leq 0.05$). The total number of elements tested at each stage is indicated below each plot.

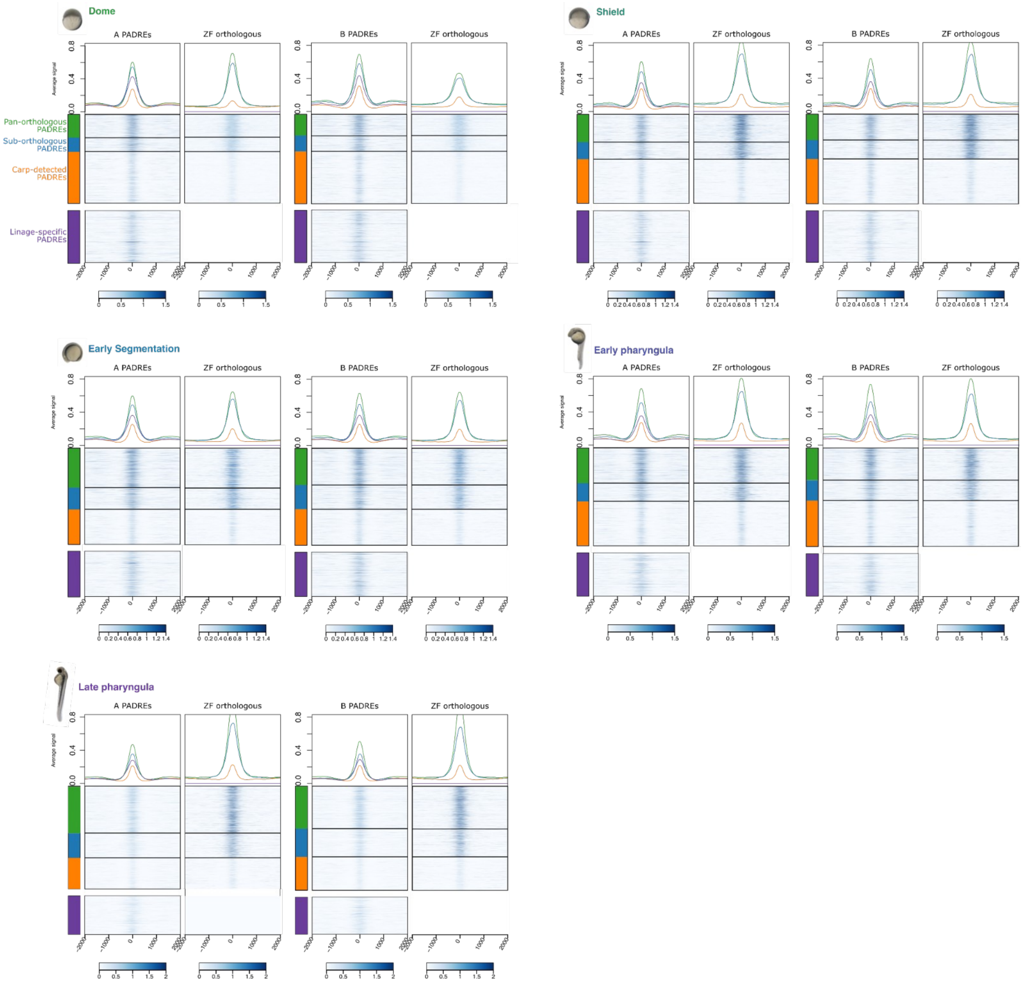


**Supplementary Figure 12. ATAC-seq chromatin accessibility profiles at carp PADREs and their projections in zebrafish across developmental stages.** Heatmaps of ATAC-seq signal centred at carp subgenome A PADREs (left pair) and subgenome B PADREs (right pair), showing the observed carp ATAC-seq signal alongside the signal at the projected coordinates in the zebrafish genome. PADREs are grouped by cross-species conservation category: pan-orthologous PADREs (green), sub-orthologous PADREs (blue), carp-detected PADREs (orange), and lineage-specific PADREs (purple). Aggregate signal profiles are shown above each heatmap. Results are displayed for five developmental stages: Dome, Shield, Early Segmentation, Early Pharyngula, and Late Pharyngula.

a

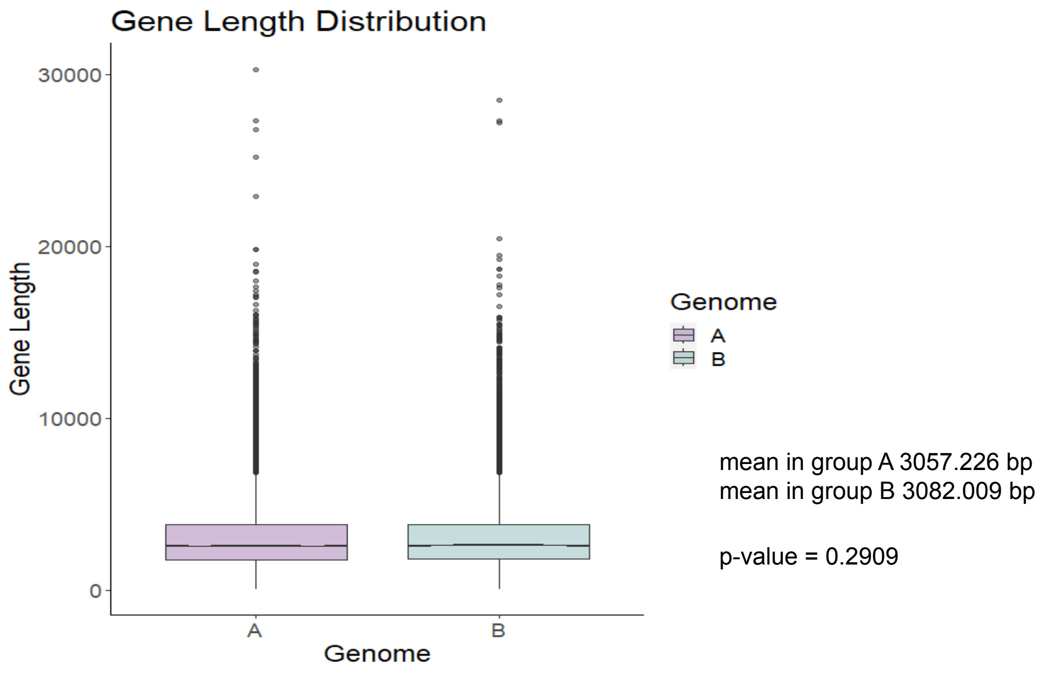


b

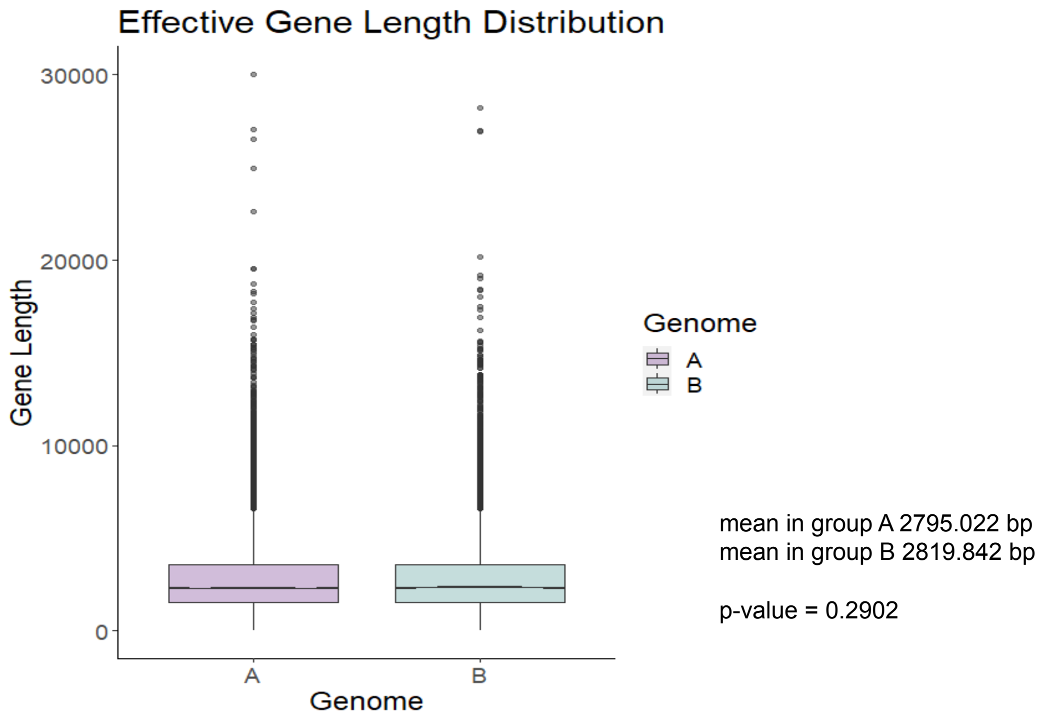


**Supplementary figure 13. Comparison of gene length distributions between subgenomes A and B. a,** Box plots showing the distribution of gene lengths (bp) for genes assigned to genome A and genome B. Mean gene lengths and the p-value from a statistical comparison between the two groups are indicated to the right of the plot. **b,** Box plots showing the distribution of effective gene lengths for genes assigned to genome A and genome B. Mean effective gene lengths and the associated p-value are indicated to the right of the plot.
